# A Self-Immobilizing NIR Probe for Non-invasive Imaging of Senescence

**DOI:** 10.1101/2020.03.27.010827

**Authors:** Jun Liu, Xiaowei Ma, Chao Cui, Ying Wang, Philip R. Deenik, Lina Cui

**Author notes:** Correspondence and requests for materials should be addressed to L.C. These authors contributed equally to this work (Jun Liu, Xiaowei Ma).

## Abstract

Cellular senescence, a process that arrests the cell cycle, is a cellular stress response to various stimuli and is implicated in aging and age-related diseases. However, the understanding of senescence in living organisms is insufficient, largely due to the scarcity of sensitive tools for the detection of cellular senescence *in vivo*. Herein, we describe the development of a self-immobilizing near-infrared (NIR) probe that can be activated by senescence-associated *β-* Galactosidase (SA-*β-*Gal), a widely accepted senescence marker. The NIR fluorophore is turned on in the presence of SA-*β-*Gal, and the self-immobilizing group, based on quinone methide chemistry, retains the fluorescence signal to the site of activation. This strategy significantly improves the sensitivity of the probe from the one we developed before. We demonstrate the non-invasive imaging of drug-induced senescence in mice models.

## Introduction

Cellular senescence, a process that halts cell proliferation, acts as an endogenous tumor suppression mechanism and is a cellular response to various stresses, including DNA damage, chromatin perturbation, and activation of oncogenes^[1]^. Increasing evidence has revealed that senescence is implicated in aging and age-related diseases^[1c, 1d]^. Senescence is regularly characterized by morphological changes *in vitro* and the overexpression of cell cycle inhibitors, such as p16, p21, and p53,^[1b, 2]^ as well as senescence-associated *β-*Galactosidase (SA-*β-*Gal), which is derived from the increased lysosomal content of senescent cells^[2-3]^. Monitoring the status of cellular senescence in living subjects allows the study of senescence in real-time without the need to terminate the experiments, enabling long-term study of senescence-related disease progression, and evaluation of treatment responses of both cancer therapies and senolytic therapies^[4]^.

Senescence-associated *β-*galactosidase (SA-*β-*Gal)^[5]^ has been widely used as a marker for senescence, and the detection of SA-*β-*Gal is mostly achieved with a colorimetric assay using 5-bromo-4-chloro-3-indoyl *β-*D-galactopyranoside (X-gal) as a chromogenic substrate^[6], [7]^, however, this approach is limited to cells and tissue sections due to its dependence on chromogenic changes^[3]^. Fluorescent probes^[8]^ developed for *β-gal* detection in *lacZ(+)* cells can potentially be used for senescence detection, and some^[9]^ have been applied for the detection of SA-*β-*Gal *in vitro*. However, these probes lack the capability of visualizing senescent cells in living animals due to short-wavelength excitation or emission of fluorophores^[5]^.

NIR fluorescent probes offer high penetration depth, minimal photodamage to tissues, and decreased background autofluorescence, and have been applied in noninvasive detection and imaging of biological targets *in vivo*^[10]^. We have previously developed a fluorogenic near-infrared (NIR) molecular probe **NIR-BG** and applied it in the imaging of drug-induced cellular senescence in different human xenograft tumor models^[11]^. We envisioned that a self-immobilizing group that could be activated by the target enzyme can further enhance the imaging efficiency by retaining the probe to the site of activation while decreasing the rapid diffusion of the freed small molecule probe^[12]^. Quinone methide chemistry has been successfully used in the design of covalent inhibitors^[13]^, immobilization of coumarin or rhodol tag ^[14]^, and photo-controlled chemical cross-linking of proteins^[15]^.

Herein, we incorporated a difluoromethyl group in the NIR fluorescent probe in which the hemicyanine skeleton is utilized as a NIR chromophore^[16]^ and a *β-*galactose residue is utilized as an enzyme recognizable trigger^[17]^, so that the electrophilic quinone methide species would be released upon activation by the SA-*β-*Gal, then trapped by the target enzymes or the nearby proteins to form a covalent linkage resulting in retained NIR signals (Figure 1). In this paper, we demonstrated the first self-immobilizing turn-on NIR fluorescent probe, NIR-BG2, for real-time imaging of cellular senescence *in vivo*.

**Figure 1.**
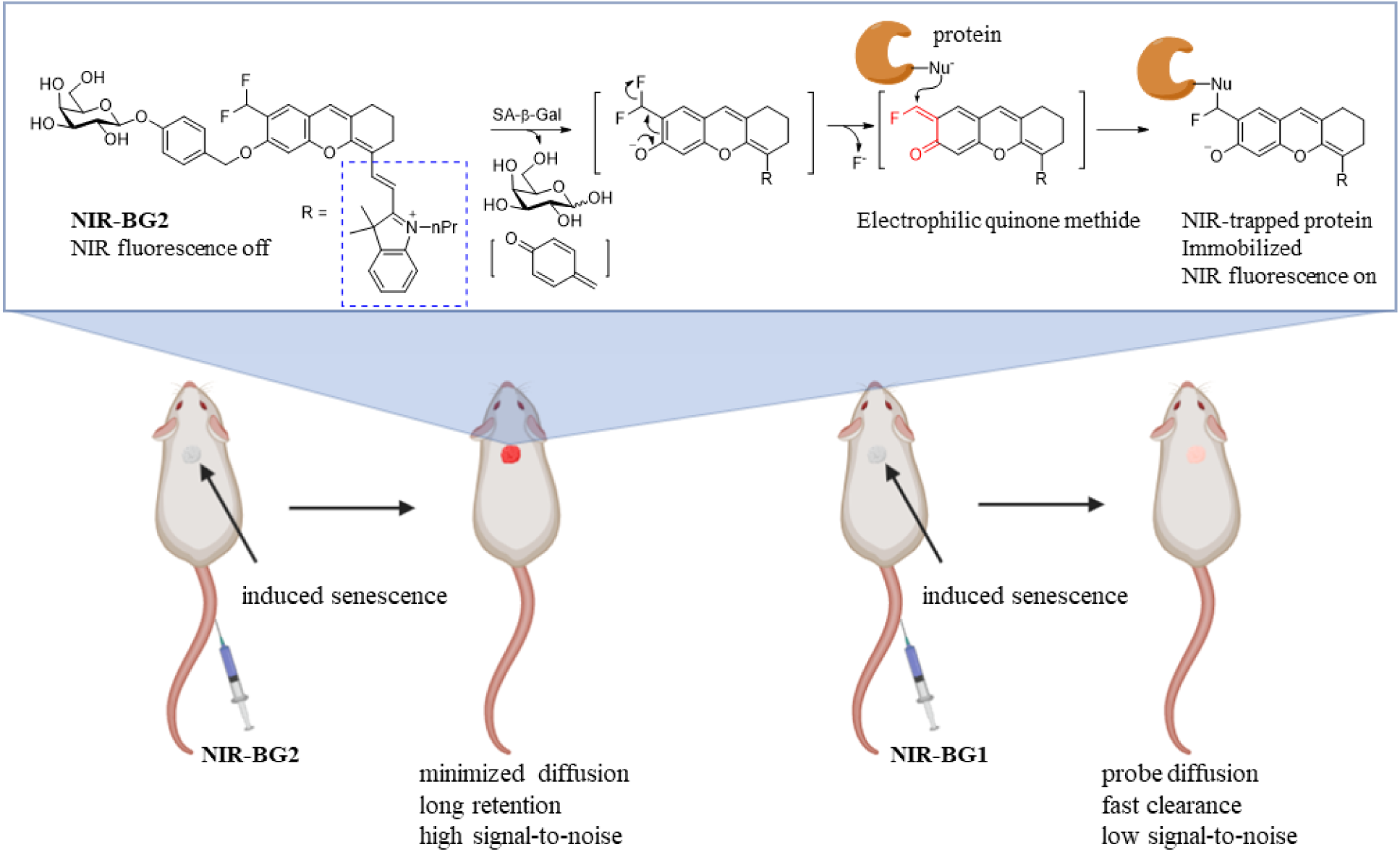
Response mechanism of self-immobilizing probes. When NIR-BG2 is hydrolyzed by the enzyme, SA-β-galactosidase, which is induced in cellular senescence upon treatment with chemotherapy reagent, NIR fluorescence is activated with long retention because the self-immobilizing group generates an electrophilic quinone methide intermediate, which is trapped by intracellular proteins simultaneously.

## Results and discussion

### Probe Design and Synthesis

It remains challenging to develop self-immobilizing NIR probes that can be turned on, as there is still a lack of efficient chemistry to install self-immobilizing functional groups under conditions that are compatible with NIR probes. In addition, even if the self-immobilizing fluorescent probe can be synthesized, the introduction of a self-immobilizing group may destabilize the fluorescent probe. For example, the QCy7-based probes ^[18]^ bearing a self-immobilizing group was found to readily decompose^[19]^, such as -CHF_2_/-CH_2_F at the ortho position of an optically tunable hydroxyl group in NIR fluorophore. In our study, (*E*)-2-(2-(6-hydroxy-2,3-dihydro-1H-xanthen-4-yl)vinyl)-3,3-dimethyl-1-propyl-3H-indol-1-ium (HXPI) was employed as the NIR fluorophore due to its high quantum yield, photostability^[20]^, and membrane permeability^[16, 21]^. More importantly, HXPI was compatible with the chemistry for the incorporation of a quinonemethide-based self-immobilizing group, and the final probe NIR-BG2 was very stable (Figure 1). The probe NIR-BG2 consists of four moieties: a *β-*Gal triggered moiety, a NIR fluorophore reporter, a self-immolative linker and a self-immobilizing moiety. To demonstrate the self-immobilizing characteristics, a control probe NIR-BG1, without the self-immobilizing properties, was synthesized as well (Figure S1). The compounds were fully characterized by mass spectrometry, ^1^H, and ^13^C NMR (and Supporting information).

### Spectroscopic Properties

We first investigated the spectroscopic properties of these two probes in PBS buffer with or without pure *β-Gal*. As shown in Figure 2, probe NIR-BG1 (5 µM) exhibited typical absorption maximum of caged HXPI at 601 nm and 650 nm; the absorption maximum for probe NIR-BG2 appeared at 596 nm and 644 nm. Upon the treatment of *β-gal*, a remarkable bathochromic shift for both probes was observed. As expected, prior to β - galactosidase treatment, both of probes NIR-BG2 and NIR-BG1 were almost nonfluorescent because the hydroxyl group of HXPI was caged with *β-*Galactosidase-triggered moiety, rendering the intramolecular charge transfer (ICT) process suppressed. However, upon addition of *β-Gal*, probe NIR-BG1 produced a dramatic fluorescence enhancement (100 fold) over the background at 699 nm which can be attributed to the enzyme-triggered cleavage of glycosylic bond to liberate the free hydroxyl group of NIR chromophore as a strong electron donor in the D-*π*-A system, thereby recovering ICT process and lighting up fluorescence^[17]^. Whereas, NIR-BG2 exhibited a small fluorescence response (16 fold) at 709 nm to *β-Gal* due to the formation of formyl group partially quenching the fluorescence ^[14b, 14d]^. To confirm the enzymatic hydrolysis mechanism, HPLC equipped with PDA detector and ESI mass spectrometry were utilized to analyze the enzymatic hydrolysis product of probes with *β-Gal*. As shown in Figure S2, a new peak was observed at 15.84 min, an indicative of NIR chromophore, in HPLC trace after incubation of 5 µM NIR-BG1 (15.23 min) with β - galactosidase (2 U) for 30 minutes. For the reaction of NIR-BG2 with β-galactosidase, the corresponding peak can be found at 16.51 min. Also, the UV-Vis spectrum of these peaks recorded by the PDA detector is in agreement with that of respective NIR chromophores. Furthermore, these peaks were subjected to ESI mass analysis and a signal was observed at m/z 412.2284 [M]^+^ for NIR-BG1, 440.2295 [M]^+^ for NIR-BG2, respectively. These results unambiguously confirmed the release of the NIR chromophore, resulting in a fluorescent response.

**Figure 2.**
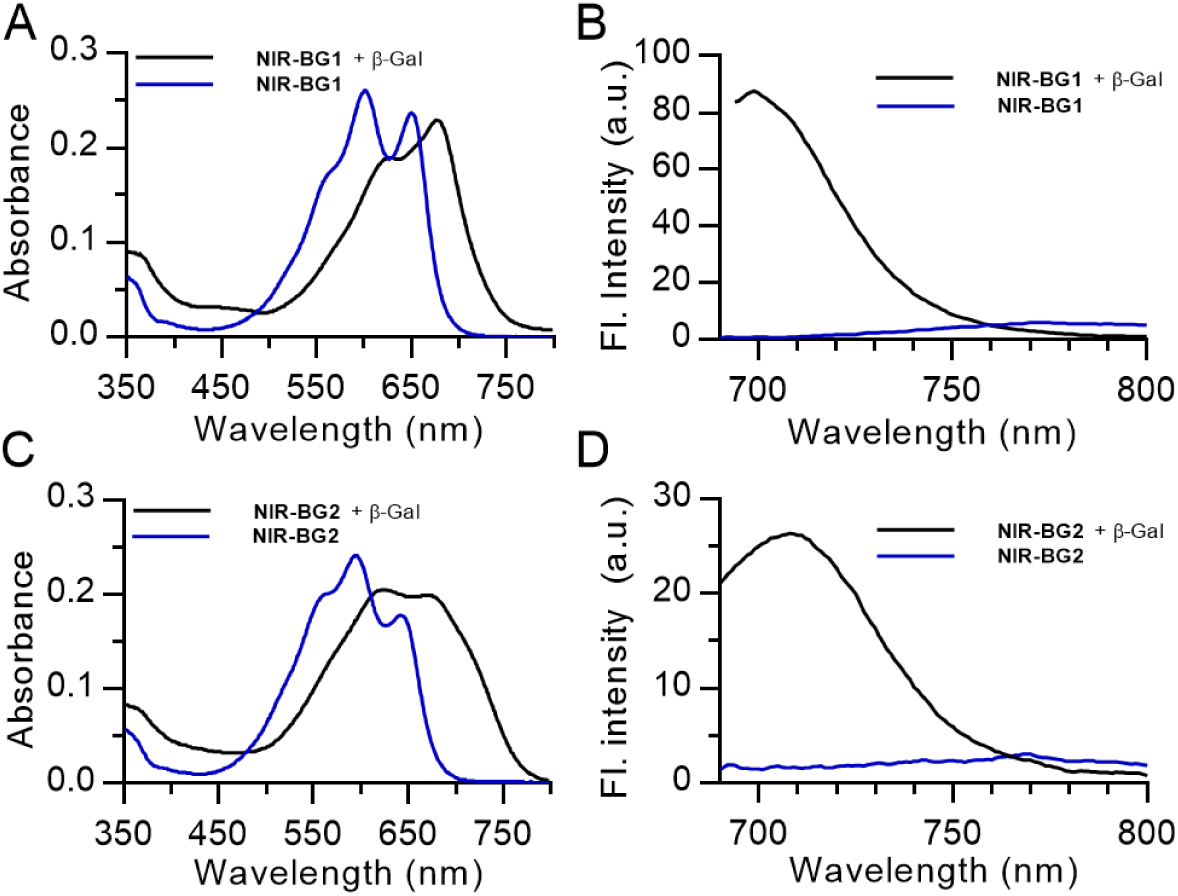
Photophysical properties of the NIR probes. UV/Vis absorption and fluorescence spectra of probes NIR-BG1 (**A, B**) and NIR-BG2 (**C, D**) (5 µM) before (blue line) and after (black line) incubation with β-Gal (2 U) in PBS (pH 7.4) buffer for 5 min at 37 °C.

### Kinetic parameters

Next, we conducted the Fluorescence-titration experiments with NIR-BG1/NIR-BG2 and various concentrations of pure *β-Gal* (0.005-0.2 U/mL). As shown in Figure S3, the fluorescence intensity dramatically increased using a high concentration of *β-Gal* in both cases. Also, the fluorescence intensity against concentrations of *β-*Galactosidase from 0.006 to 0.2 U/mL exhibited a good relationship at 700 nm for NIR-BG1, 708 nm for NIR-BG2, respectively. The regression equation is calculated as F_700 nm_ = 324.5[*β-gal*] +1.671 for NIR-BG1 and F_708 nm_ = 554.9[*β-gal*] + 0.452 with a linear coefficient of 0.9957 and 0.9972 respectively. The kinetic parameters such as the Michaelis constant (*K*_m_), the turnover number (*k*_cat_), and the catalytic efficiency constant (*k*_cat_/*K*_m_), were subsequently studied by monitoring the fluorescent intensity change at various concentrations of probes with β-galactosidase (0.1 U/mL) to obtain the enzymatic hydrolysis reaction rate (Figure S4). Then the kinetic parameters were determined by plotting the Lineweaver-Burk equation: 1/V_0_ = *K*_m_/*k*_cat_[E_0_][S] + 1/*k*_cat_[E_0_], where [E_0_] is the concentration of *β-gal*. Thus, the kinetic parameters *K*_m_, *k*_cat_ and *k*_cat_ / *K*_m_ were calculated to be 2.0 μM, 6.4 s^-1^, and 3.2 μM^-1^•s^-1^ for NIR-BG1; 9.3 μM, 14.6 s^-1^, and 1.6 μM^-1^•s^-1^ for NIR-BG2.

### Fluorescence Western blot analysis

To validate the self-immobilization of probe NIR-BG2, the fluorescence western blot analysis was carried out as it enables the quantification of *β-gal* and probe concentration under different fluorescence channels to observe the colocalization of NIR-BG2 and *β-gal* enzyme to verify the activation and self-immobilization of the probe. Also, a quantitative analysis could be performed to estimate the relationship between the *β-gal* concentration and the probe activation and binding. The results showed that the bright NIR-BG2 fluorescence signal exactly colocalized with the signal from the *β-gal* enzyme while NIR-BG1 signal was very weak (Figure 3A, Figure S5), which demonstrated the NIR-BG2 could be activated and bind to the *β-gal* enzyme but NIR-BG1 cannot. The quantification data indicated the linear correlation between the concentration of *β-gal* enzyme and the fluorescence intensity of NIR-BG2, which demonstrated the ability of NIR-BG2 for quantitatively imaging the cellular senescence by measuring the concentration of *β-gal* enzyme both *in vitro* and *in vivo* (Figure 3B, 3C).

**Figure 3.**
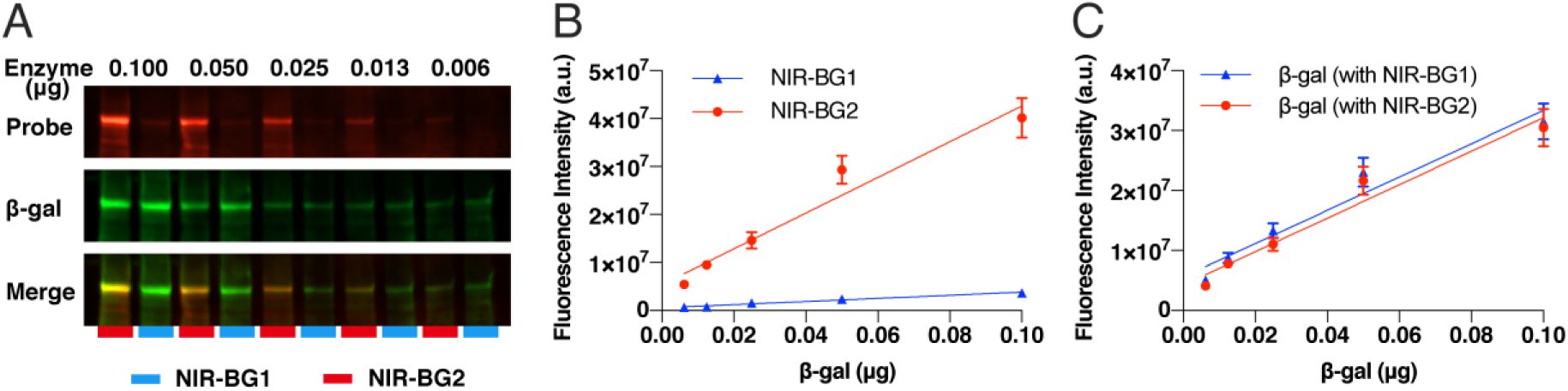
Fluorescent western blot of NIR-BG1 and NIR-BG2 (5 μM) incubated with different concentrations of recombinant β-galactosidase for 4 hours. (**A**) Fluorescence western blot imaging. (**B**) the fluorescence intensity of the probes. (**C**) The fluorescence intensity of β-galactosidase.

### Flow cytometry

To quantitatively estimate the cell uptake and activation of NIR-BG1 and NIR-BG2 in therapy-induced senescent cancer cells, flow cytometry analysis was performed. It showed significant higher fluorescence intensity in senescent HeLa cells and CT26.CL25 cells compared to the untreated HeLa cells (Figure 4, Figure S6) and CT26.WT cells (Figure S7). Meanwhile, the fluorescence signal from NIR-BG2 was significantly higher than that from NIR-BG1 in senescent HeLa cells, which demonstrated the continuous activation and accumulation of NIR-BG2 due to the self-immobilizing effect. Furthermore, the fluorescence intensity increased following the incubation time. These results further certified the ability of the NIR-BG2 to monitor cellular senescent by targeting the SA-*β-*Gal activity.

**Figure 4.**
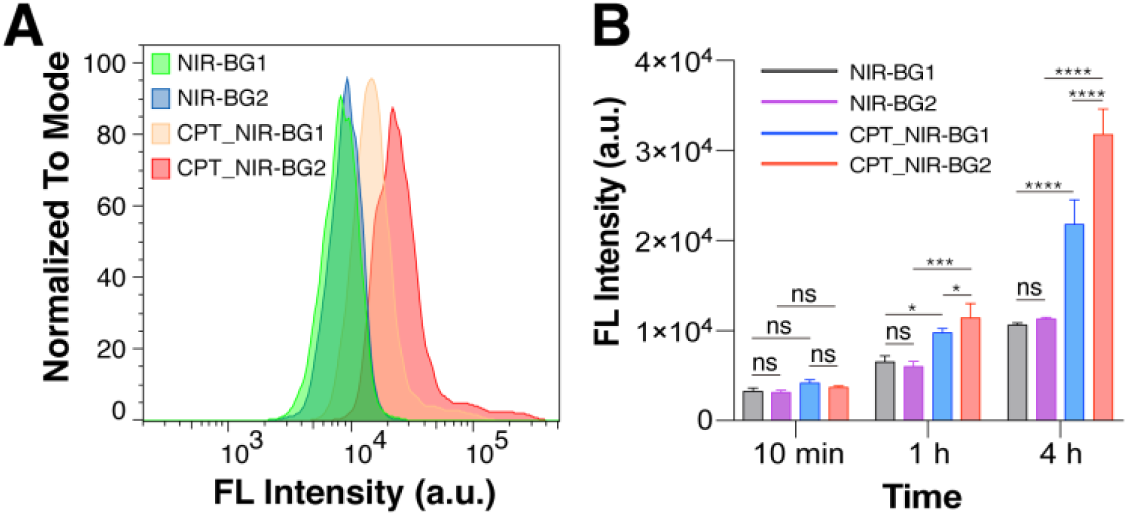
(**A**) Flow Cytometric analyses of untreated and CPT-treated HeLa cells incubated with NIR-BG1 or NIR-BG2 for 4 h. (**B**) Quantitative analysis of flow cytometry showed that the cellular uptake of the probes in CPT induced senescent HeLa cells are higher than healthy HeLa cells, but NIR-BG2 is better than NIR-BG1. (λex/λem = 642 nm/675±25 nm) (*\* p<0*.*05, *** p<0*.*0005, **** p<0*.*0001*).

### Immunofluorescence cell staining

To investigate the imaging ability and the difference between NIR-BG2 and NIR-BG1 to detect SA-*β-*Gal in senescent cells, HeLa cells were imaged after inducing senescence and compared with untreated HeLa cells. It could be observed that the CPT-treaded HeLa cells are significantly enlarged than untreated HeLa cells, indicating the morphology change of the senescence cells. Although NIR-BG1 and NIR-BG2 showed fluorescence signals in the senescent HeLa cells and highly colocalized with the SA-*β-*Gal staining, the signal of NIR-BG2 was significantly higher than that of NIR-BG1 (Figure 5).

**Figure 5.**
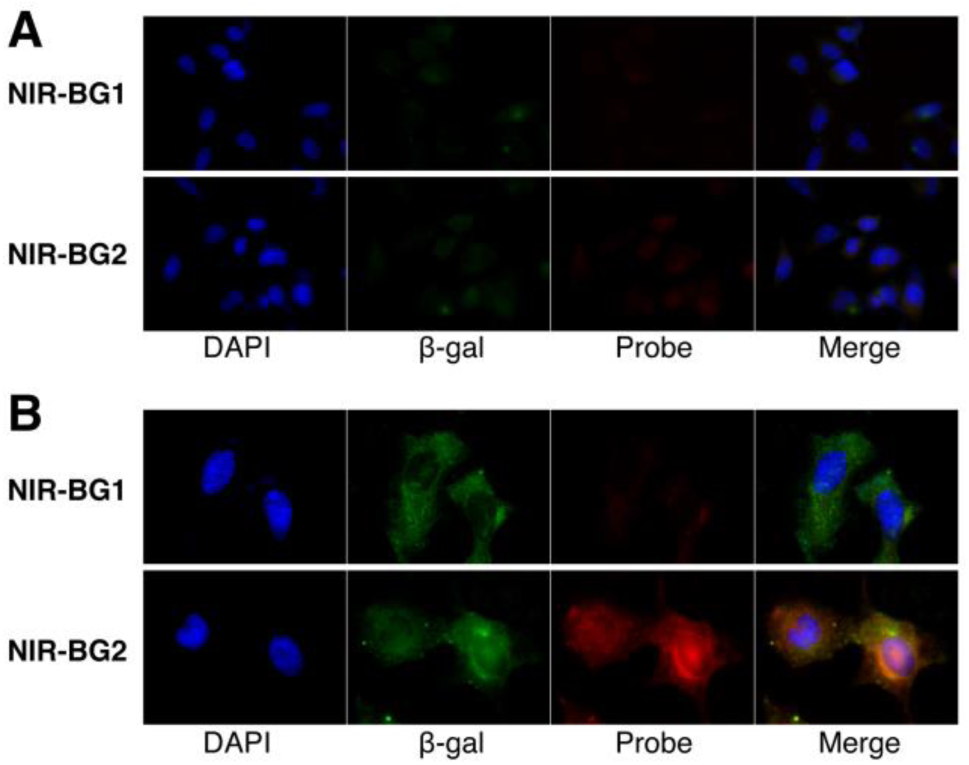
Fluorescent microscope images of (**A**) untreated and (**B**) CPT-treated HeLa cells incubated with NIR-BG1 or NIR-BG2 (5 μM) for 2 hours and followed by incubation with fresh medium for another 4 hours. (λex/λem = 395 nm/460 nm for DAPI, 470 nm/535 nm for β-actin and 740 nm/767 nm for probe) scale bar 20 um.

The dynamic clearance of probes in CT26.CL25 cells showed that the NIR-BG1 would be cleared out within 24 h while the NIR-BG2 could accumulate in the cells for more than 24 hours (Figure S8). These results indicated that the higher fluorescence in senescent cells was due to the attachment of activated NIR-BG2 to the SA-*β-*Gal and some other proteins. Thus, it demonstrated that the self-immobilizing NIR-BG2 is more specific and sensitive to detect the *β-gal* in senescent cells and would be used for further *in vivo* imaging of chemotherapy-induced cancer senescence.

### *In vivo* animal imaging

*In vivo* fluorescence imaging of HeLa xenografts was successfully performed to evaluate the capability of NIR-BG1 and NIR-BG2 to visualize chemotherapy-induced cancer senescence *in vivo*. HeLa tumor-bearing mice were treated with CPT to induce the cancer senescence following a previously reported method by our group. The CPT treated tumor showed significantly increased fluorescence intensity compared to the saline-treated tumor (Figure 6A). The quantification of the optical imaging (Figure 6B) showed that the fluorescence intensity at 24 h from the CPT treated tumor was (1.04 ± 0.15) × 10^7^ and (0.53 ± 0.13) × 10^7^ (NIR-BG2 *vs* NIR-BG1). Importantly, compared with NIR-BG1, the NIR-BG2 showed much higher signal and longtime accumulation in the CPT treated tumors due to the attachment of it to the proteins after being activated by *β-gal*. The *ex vivo* imaging (Figure 7A) and quantitative (Figure 7B) analysis of the tumors and organs further confirmed the *in vivo* imaging results.

**Figure 6.**
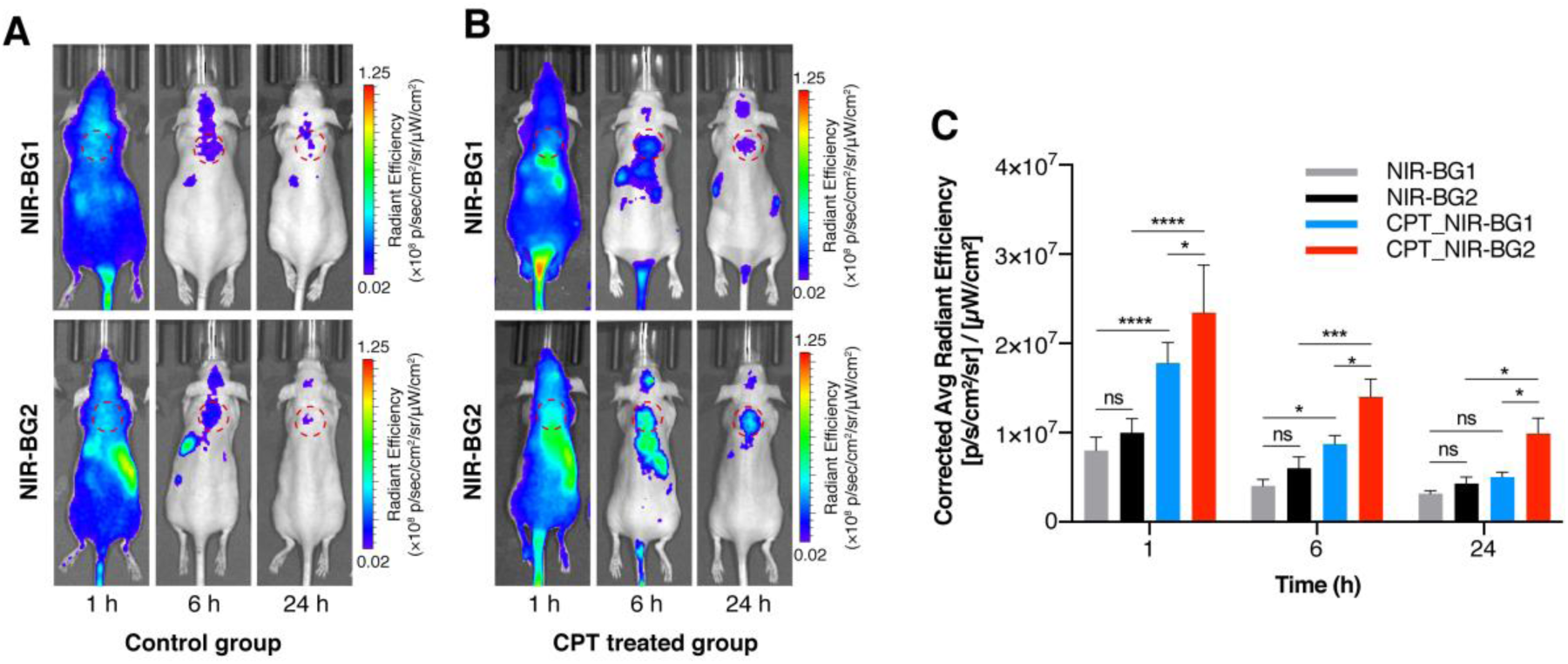
Animal imaging of (**A**) saline-treated (control) and (**B**) CPT-treated HeLa xenografts mice after intravenous tail injection of NIR-BG1(top) or NIR-BG2 (bottom) at 1 h (left), 6 h (middle) and 24 h (right). (**C**) The quantitative analysis of the *in vivo* fluorescence imaging showed that the uptake of NIR-BG2 in senescent tumors is significantly higher than NIR-BG1 at different time points after injection. (λex/λem = 675 nm/720 ± 20 nm) (*\* p<0*.*05, *** p<0*.*0005, **** p<0*.*0001*).

**Figure 7.**
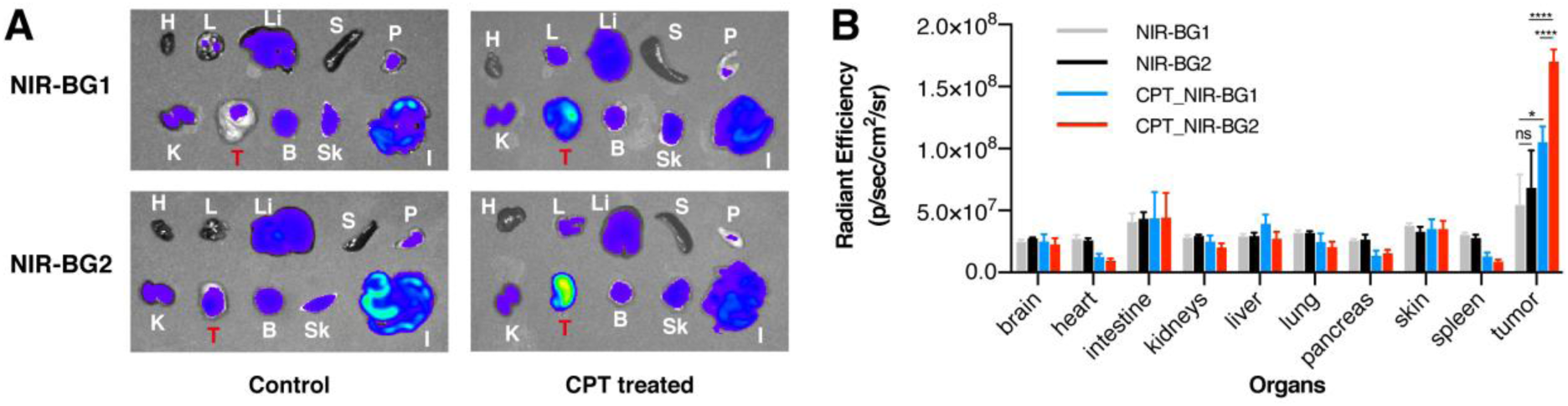
*Ex vivo* imaging (**A**) and quantification (**B**) of tumors and major organs of control and CPT treated HeLa xenografts. (H = heart, L = lung, Li = liver, S = spleen, P = pancreas, K = kidneys, T = tumor, B = brain, Sk = skin, I = intestine) (λex/λem = 675 nm/720 ± 20 nm) (*\* p<0*.*05, **** p<0*.*0001*).

X-gal staining and fluorescence imaging of the HeLa tumor slices confirmed the probes are accumulated in the chemotherapy-induced senescent cancer cells and (Figure 8), and also certified the NIR-BG2 is better than NIR-BG1 for detecting senescent tumor cells *in vivo*.

**Figure 8.**
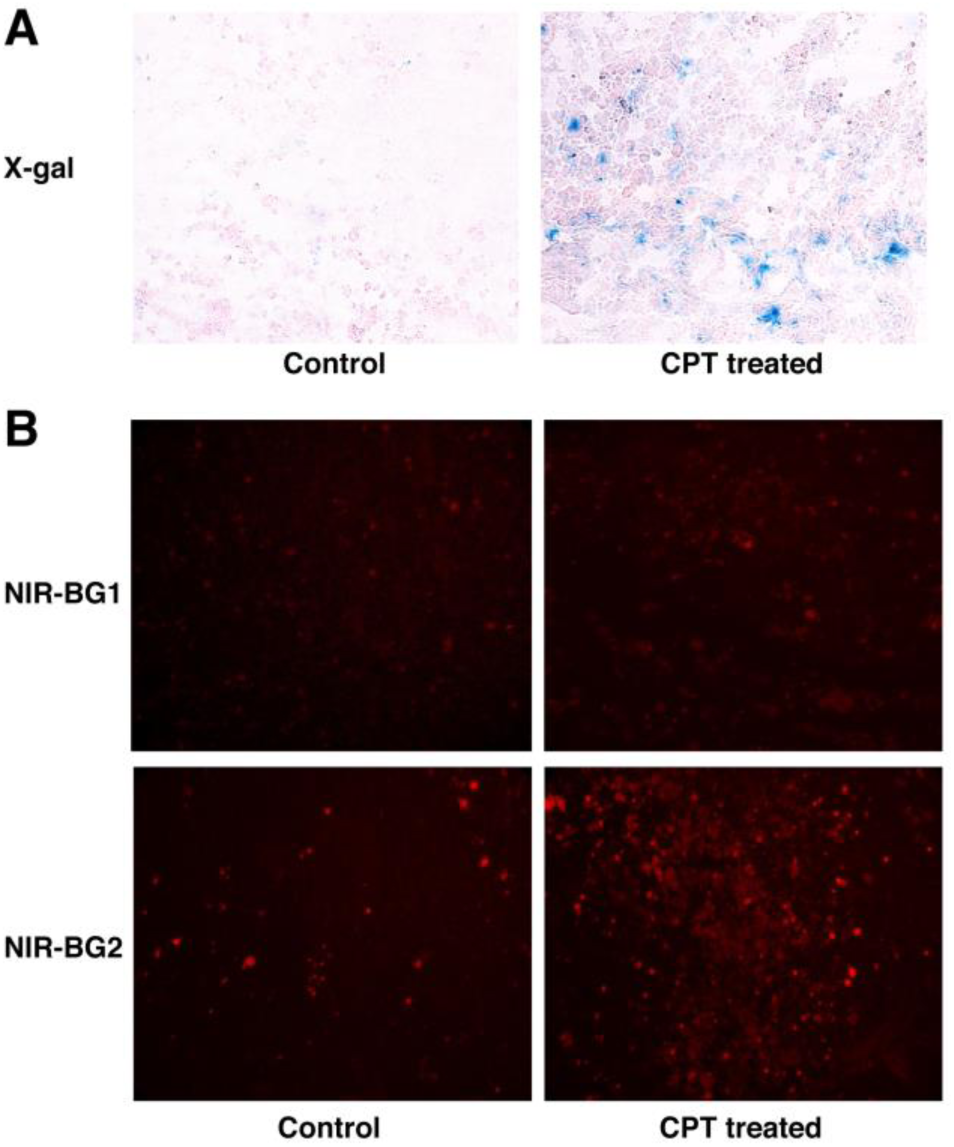
Pathological staining of tumor sections. (**A**) Conventional X-Gal (blue) staining for SA-*β-* Gal activity and Eosin (pink) staining of HeLa tumor slides without (control) or with CPT treatment. (**B**) Fluorescent imaging of control (left) and CPT-treated (right) HeLa tumor sections with NIR-BG1 (top) or NIR-BG2 (bottom) (λex/λem = 675 nm/720 ± 20 nm).

## Conclusion

We have developed a self-immobilizing NIR probe for the imaging of cellular senescence in living animals. This self-immobilizing NIR was efficiently activated by SA-*β-*Gal to produce intense fluorescence signals both *in vitro* and *in vivo*. Compared with the control probe **NIR-BG1** lacking the immobilizing group, the probe **NIR-BG2** showed stronger NIR fluorescence after activation in the senescent cells and longer retention inside the senescent cells. Importantly, **NIR-BG2** showed significant retention in animal models induced with senescence, offering a wider time window to allow the clearance of background fluorescence signals in circulation. We anticipate that the combined advantage of the NIR optical probe and long-term tracking makes our self-immobilizing probe useful for investigating the senescence in a living organism.

## Materials and methods

### Materials

4’, 6-diamidino-2-phenylindole (DAPI) was purchased from Biotium, CA, USA. Bovine serum albumin (BSA), Fetal Bovine Serum (FBS), and PBS were obtained from VWR, PA, USA. Dulbecco’s Modified Eagle’s medium (DMEM) and Eagle’s minimal essential medium (EMEM) were from Corning Inc, USA. *β-*Galactosidase (*E. coli*) was purchased from Abnova (catalog #P5269). Goat anti-rabbit IgG H&L-AF488 (Catalog #ab150077) was purchased from Abcam (USA). Anti-*β-*Galactosidase antibody (Catalog # A-11132), and goat anti-rabbit IgG H&L Secondary Antibody, HRP (Catalog # 65-6120), ActinGreen 488 ReadyProbes (Catalog # R37110) and ProLong Gold Antifade Mountant were purchased from Invitrogen, USA. Radioimmunoprecipitation assay (RIPA) cell lysis buffer was from Enzo Life Sciences, NY, USA. HeLa (human cervical cancer cell line), CT26.WT (wild type mouse colon fibroblast carcinoma cells) and CT26.CL25 (*lacZ+* CT26 cell, engineered cells that highly express *β-gal*) cell lines were purchased from American Type Culture Collection (ATCC), VA, USA. Mini-PROTEAN^®^ TGX™ Precast Gels and polyvinylidene fluoride (PVDF) membrane were purchased from Bio-Rad, USA. Dimethyl sulfoxide (DMSO) was from Sigma-Aldrich, USA.

### Cells culture and induction of cellular senescence

HeLa cells were cultured at 37 °C in EMEM supplemented with 10% FBS and 1% penicillin under 5% CO_2_ and 95% humidity. CT26.CL25 and CT26.WT cells were cultured in complete DMEM medium supplemented with 10% FBS and 1% penicillin at 37 °C with 5% CO_2_ and 95% humidity. To induce cellular senescence, HeLa cells were cultured with freshly prepared culture medium containing 7.5 nM CPT and 5% DMSO for 7 days.

### Fluorescence Western blot analysis

To verify the binding affinity of activated NIR-BG1 and NIR-BG2, different concentration of the *β-gal* enzyme was incubated with 5 μM probes (in 1× PBS containing 5% DMSO) for 4 hours at 37 °C. Then samples were separated using 4-20% Mini-PROTEAN^®^ TGX™ Precast Gels. After being transferred onto the PVDF membrane, *β-gal* was stained with anti-*β-*Galactosidase antibody (1:5000) and goat anti-chicken IgY H&L-AF568 (1:5000). Then the PVDF membrane was imaged using ChemiDoc MP (Bio-Rad, USA) to observe the co-localization of *β-gal* bands and probe signals.

### Immunofluorescence cell staining

HeLa cells were seeded on glass coverslips (0.13-0.16 mm thickness) at a density of 3 × 10^4^ cells/well (total 8 wells, 4 wells for inducing senescence and 4 wells for control) and cultured overnight. At the next day, cells were treated with CPT (20 nM) or PBS for 4 days. After treatment, all cells were incubated with 5 μM probe (NIR-BG1 or NIR-BG2) for 2 hours, Then the culture medium was changed to probe-free MEM to allow cells to wash out the probes for 0 and 4 hours. Cells were then fixed and subsequently stained with actin (ActinGreen 488 ready probes reagent 2 drops/ml) for 15 min and DAPI for 5min. Fluorescence microscopy imaging was performed to observe and compare the residual of probes in cells. Then, all coverslips were mounted on the slides with ProLong Gold Antifade Mounting media and were acquired for fluorescent microscope imaging (Nikon Ti2, Japan). CT26.WT and CT26.CL25 cells were also stained and image with same procedure just without treatment to confirm the uptake and binding of the probes.

Dynamic clearance of the probes in the *β-gal* overexpressed CT26.CL25 cells was also performed to estimate the pharmaceutical kinetics of the probes in cells. CT26.CL25 cells were incubated with probes (5 μM) for 2 hours. Then the culture medium was changed to probe-free DMEM to allow cells to wash out the probes for 0, 1, 4, and 24 hours. Cells were then fixed and subsequently stained with *β-*actin and DAPI. Fluorescence microscopy imaging was performed to observe and compare the residual of probes in cells.

### Flow cytometry

HeLa cells were seeded in a 24-well plate at a density of 3 × 10^5^ cells/well with or without CPT treatment (20 nM) for 7 days. CT26.WT and CT26.CL25 cells were seed in a 24-well plate at a density of 4 × 10^5^ cells/well and cultured at 37 °C overnight. Then cells were incubated with 5 μM of probes for 10 min, 1 hour, and 4 hours at 37 °C. After being washed 3 times with PBS, cells were digested with 0.25% trypsin and resuspended in 200 μL PBS buffer for flow cytometry analysis (Accuri C6 Plus, Becton Dickinson and Company, USA).

### In vivo imaging

*In vivo* imaging was performed in accordance with protocols approved by the Institutional Animal Care and Use Committee of the University of New Mexico and followed the National Institutes of Health guidelines for animal care. Twenty thymic female nude mice (from Harland Laboratories) were injected subcutaneously with 2 × 10^6^ HeLa cells to establish a tumor xenograft model. When tumor size reaches 100 mm^3^, mice were randomly divided into 4 groups for treatment. Two groups were administrated with CPT by gavage (2 mg/kg, every two days for 4 times, total dose was 8 mg/kg) and two groups were given saline for control. At day 10, mice were injected with probes (10 nmol, 100 μL) through tail vein. The fluorescence images were acquired at 1 h, 6 h, and 24 h post-injection using an IVIS Spectrum optical imaging system (PerkinElmer, USA) with a 680 nm excitation and 720 nm emission filter set. All mice were euthanized after the last imaging for tumors and major organs collection, *ex vivo* imaging. After a fast imaging, all tumors were immersed in optimal cutting temperature (OCT) compound and frozen on dry ice immediately for later immunohistochemistry experiments.

### Immunohistochemistry Staining

The frozen tumors were sectioned into 4 μm slices and were fixed with 4% formaldehyde for 10 min. The slides were then stained with Eosin and X-gal according to our previously reported method^[11]^.

### Statistical analysis

Values are reported as the mean ± standard deviation unless otherwise noted. Student’s *t-test* and two-way analysis of variance (ANOVA) were used to determine the statistical significance with probability values less than 0.05 (*p* < 0.05). All statistical calculations were performed using Prism 7.0 (GraphPad Software).

## Acknowledgements

The work was supported by research grants to Prof. L. Cui from the University of New Mexico (UNM Startup Award), National Institute of General Medical Sciences of National Institutes of Health (Maximizing Investigators’ Research Award for Early Stage Investigators, R35GM124963), the UNM Comprehensive Cancer Center, and the National Cancer Institute of the United States (P30CA118100), and the University of Florida (UF Startup Fund). We thank Prof. Wei Wang (UNM) for the use of his fluorescence spectrometer, UNM Fluorescence Microscopy Shared Resource for the use of the confocal microscopes, and Dr. Mara Steinkamp at the UNM Cancer Center Animal Models Shared Resource for assistance in the animal studies.

## Author Contributions

J.L. and L.C. conceived the project. J.L. performed the syntheses of all compounds, measured the absorption and fluorescence spectrum, and performed kinetic studies. X.M., C.C., Y.W. and P. R. D. performed *in vitro* assays. X.M., Y.W., and C.C. performed confocal microscopy. Y.W., X.M., C.C., performed flow cytometry. X.M., C.C. performed all the *in vivo* experiments. J.L., X.M., and L.C. processed and analyzed the data, and wrote the paper. All authors commented on the manuscript.

## Additional Information

Supplementary information related to this article is available. Conflicts of Interest: The authors declare no conflict of interest.

