## Supporting information for "A Self-Immobilizing NIR Probe for Non-invasive Imaging of Senescence"

Deuterated solvents were purchased from Sigma-Aldrich and Merck Millipore. Compound **1**[1] and NIR fluorophores **2a**, **2b**[2] were prepared following a literature procedure. $^{1}H$ and $^{13}C$ NMR spectra were recorded on a Bruker instrument (500 and 126 respectively) and internally referenced to the residual solvent signals ($^{1}H$: δ 7.26; $^{13}C$: δ 77.16 for $CDCl_3$, $^{1}H$: δ 3.31; $^{13}C$: δ 49.0 for $CD_3OD$ respectively). NMR chemical shifts (δ) and the coupling constants (J) for $^{1}H$ and $^{13}C$ NMR are reported in parts per million (ppm) and in Hertz, respectively. The following conventions are used for multiplicities: s, singlet; d, doublet; t, triplet; m, multiplet; and dd, doublet of doublet. High resolution mass was recorded on Waters LCT Premier Mass Spectrometer. Absorption spectra were taken on Shimadzu UV-1800 UV-VIS Spectrophotometer. Fluorescence spectra were recorded on Edinburgh FLS980 fluorescence spectrometer and Shimadzu RF-5301pc spectrophotometer. β-Galactosidase (*E. coli.*) (catalog. P5269) was purchased from Abnova. HPLC analysis was performed on Thermo Scientific Dionex UltiMate 3000 with a reverse phase column C18 250x10 mm (Teledyne Isco).

**Experimental section**

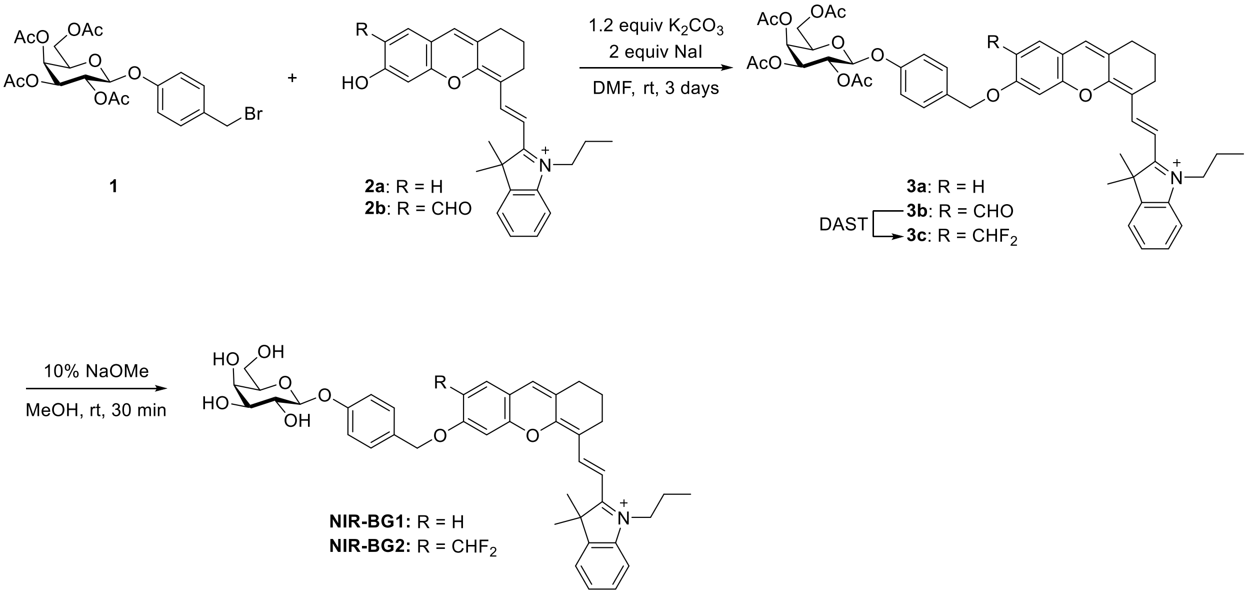

**Figure S1.** Synthetic route to the probes.

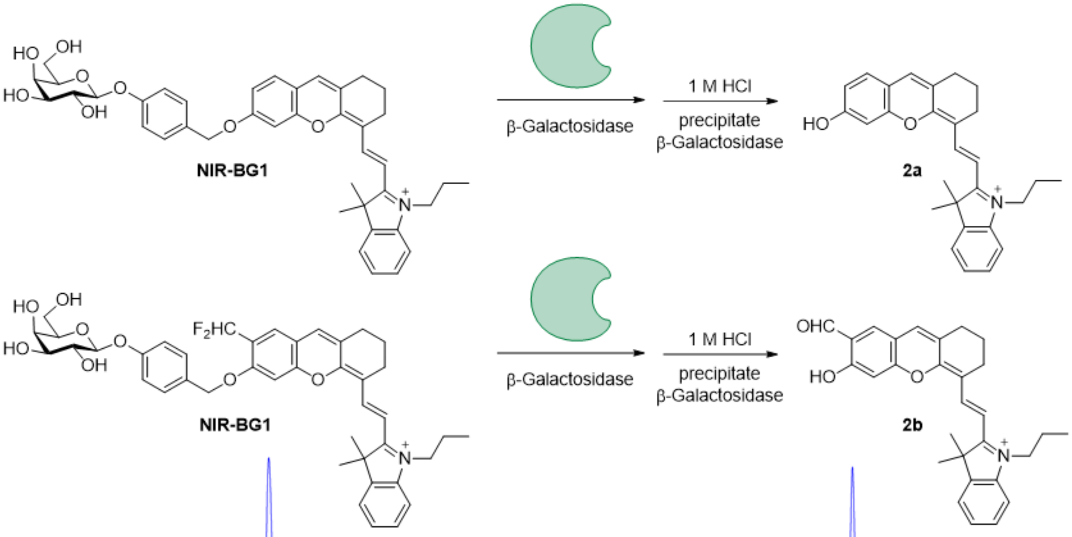


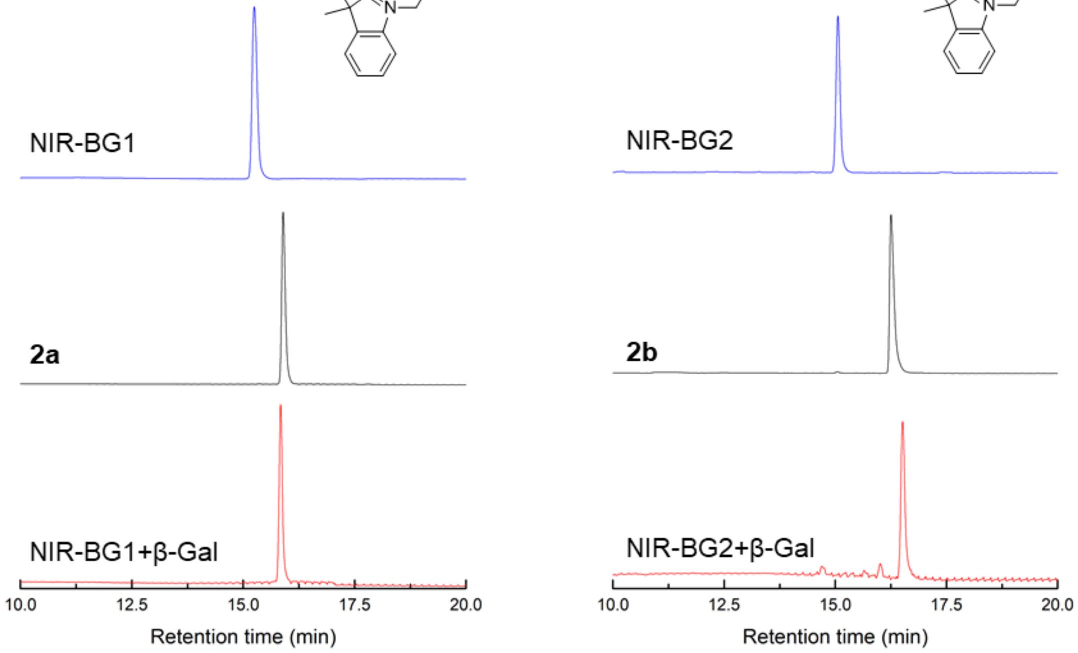


**Fig. S2.** HPLC traces of NIR-BG1 (left) and NIR-BG2 (right) probe only (top), standard sample of hydrolyzed end product (middle), NIR-BG probe with β-gal (bottom).

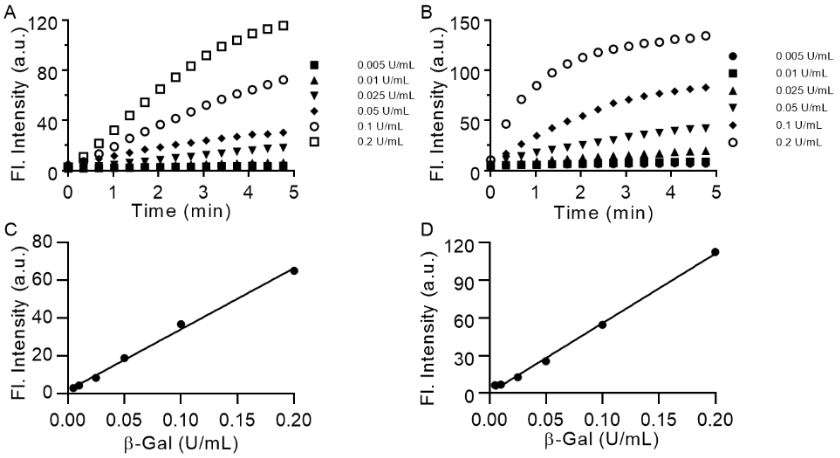


**Figure S3.** Time-dependent fluorescence intensity increment using A) NIR-BG1 (5 μM) and B) NIR-BG2 with various amounts of β-Galactosidase. C) Fluorescent intensity of C) NIR-BG1 and D) NIR-BG2 activation by various concentrations of β-Galactosidase at 2 min. λex/em = 679 nm/700 nm for NIR-BG1; λex/em = 675 nm/708 nm for NIR-BG2.

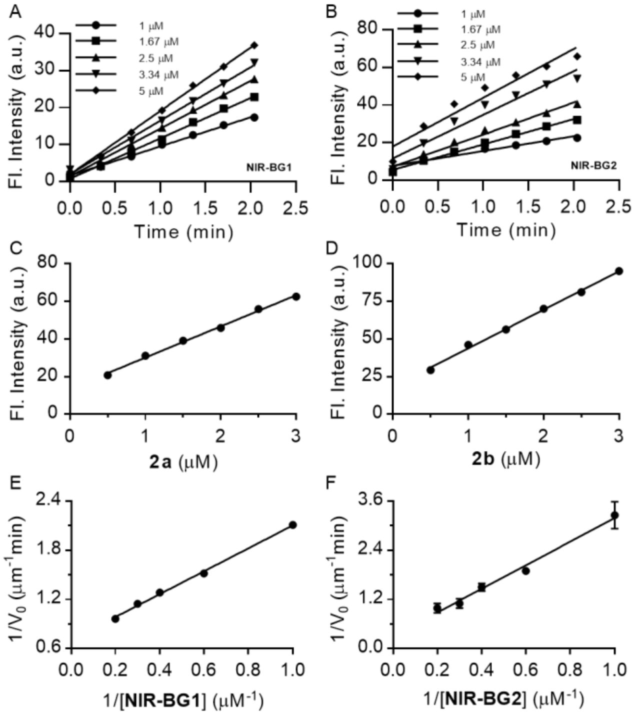

**Figure S4.** Kinetic parameters determination. Time-dependent fluorescence intensity increment using β-Galactosidase (0.1 U/mL) with different concentrations of A) NIR-BG1 and B) NIR-BG2. Standard fluorescence curve of C) **2a** and D) **2b** at different concentrations. λex/em = 679 nm/700 nm for NIR-BG1 and **2a**; λex/em = 675 nm/708 nm for NIR-BG2 and **2b**. Lineweaver-Burke plot of E) NIR-BG1 and F) NIR-BG2 activation by β-Galactosidase.

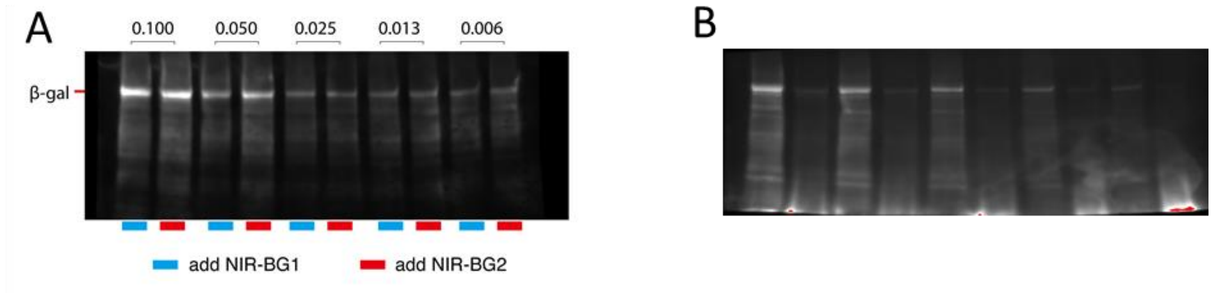


**Figure S5.** Fluorescence image of the whole membrane. A) The imaging of membrane stained with anti-β-gal antibody; B) The imaging of membrane at probe channel.

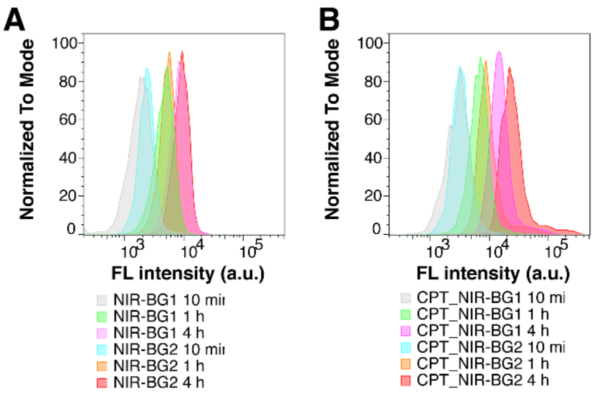


**Figure S6.** Flow Cytometry analysis of (A) untreated HeLa cells and (B) CPT treated HeLa cells incubated with NIR-BG1 or NIR-BG2 for 10 min, 1 h, and 4 h. The uptake of the probes in the CPT treated group was higher than untreated group and continuously increased over time. Furthermore, the NIR-BG2 showed higher accumulation than NIR-BG1 in CPT treated cells. (λex/λem = 642 nm/675 ± 25 nm)

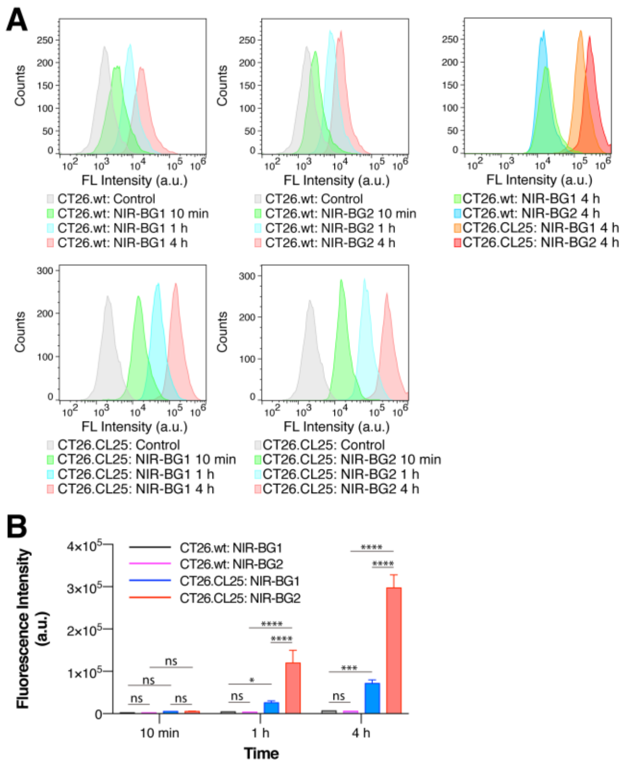


**Figure S7.** Flow Cytometry (A) and quantitative analysis (B) of *LacZ(+)* CT26 cells (CT26.CL25, overexpress β-gal) and wild type CT26 cells (CT26.wt) incubated with NIR-BG1 or NIR-BG2 for 10 min, 1 h, and 4 h. The uptake of the probes in the β-gal overexpressed CT26.CL25 cells was higher than control CT26 cells and continuously increased over time. Furthermore, the NIR-BG2 showed higher accumulation than NIR-BG1 in CT26.CL25 cells. (λex/λem = 642 nm/675 ± 25 nm) (* $p<0.05$, *** $p<0.0005$, **** $p<0.0001$).

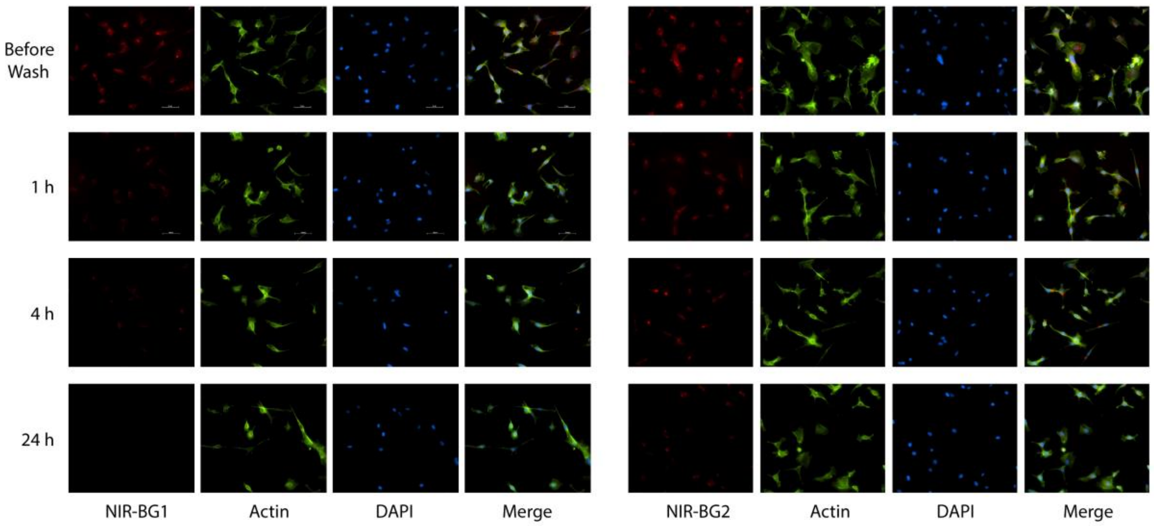


**Figure S8.** Fluorescence microscope images of CT26.CL25 cells (clearance experiment). CT26.CL25 cells were incubated with NIR-BG1 or NIR-BG2 for 4 h in 37 ºC. Then cells were washed with PBS for 3 times and the culture medium was replaced. Cells were then cultured for different periods (1 h, 4 h, and 24 h) to let the probe being cleared from cells. The NIR-BG1 was cleared out from cells after 4 hours but NIR-BG2 accumulated in cells for at least 24 hours.

### UV-Vis and fluorescence spectrum

To a solution of 20 µL of 50 µM probes NIR-BG1 or NIR-BG2 and 160 µL PBS (pH = 7.4) was added 20 µL 0.1U/µL β-Galactosidase to obtain 5 µM probes with 2 U β-Galactosidase solution. After incubation at 37 °C for 5 min, the reaction solution was transferred to quartz cuvettes to measure absorbance or fluorescence. Absorbance spectrum scan range from 350 nm to 800 nm (1 nm increment). fluorescence spectrum setting for NIR-BG1: λex = 679 nm, Slit Width 5 nm. fluorescence spectrum setting for NIR-BG2: λex = 675 nm, Slit Width 5 nm. Emission was record from 690 nm to 800 nm, Slit Width 5 nm. Absorption spectra were recordes on Shimadzu UV-2700 UV-VIS Spectrophotometer. Fluorescence spectra were recorded on Shimadzu RF-5301pc spectrophotometer.

### Time-dependent fluorescence intensity increment using β-Galactosidase (0.1 U/mL) with different concentrations of probes.

To a solution of 4, 6.7, 10, 13.3, 20 µL of 50 µM probe NIR-BG1 or NIR-BG2 and 176, 173.3, 170, 166.7, 160 µL PBS (pH = 7.4) buffers was added 20 µL β-Galactosidase (1 U/mL) in quartz cuvettes. Then final concentration of NIR-BG1 or NIR-BG2 is 1.0, 1.67, 2.5, 3.34, 5 µM and β-Galactosidase is 0.1 U/mL. Then the fluorescence intensity was recorded on Shimadzu RF-5301pc spectrophotometer at each timepoint. fluorescence spectrum setting for NIR-BG1: λex/λem = 679/700 nm, Slit Width 5 nm/ Slit Width 5 nm. Fluorescence spectrum setting for NIR-BG2: λex/λem = 675/708 nm, Slit Width 5 nm/ Slit Width 10 nm.

**Time-dependent fluorescence intensity with various amounts of β-Galactosidase**

To a solution of 20 µL of 50 µM probe NIR-BG1 or NIR-BG2 and 160 µL PBS (pH = 7.4) buffer was added 20 µL β-Galactosidase with various concentrations (0.05-2.0 U/mL) in quartz cuvettes. Then final concentration of β-Galactosidase is from 0.005-0.2 U/mL. Then the fluorescence intensity was recorded on Shimadzu RF-5301pc spectrophotometer at each timepoint. fluorescence spectrum setting for NIR-BG1: λex/λem = 679/700 nm, Slit Width 5 nm/ Slit Width 5 nm. Fluorescence spectrum setting for NIR-BG2: λex/λem = 675/708 nm, Slit Width 5 nm/ Slit Width 10 nm.

**HPLC analysis of NIR-BG1 and NIR-BG2 activation by β-galactosidase**. β-Gal (5 unit, 50 µL) was added to the NIR-BG probe solution (30 µL, 1 mM) and the mixture was incubated for 10 minutes at 37 °C. The resulting mixture was then quenched with HCl (1 M, 120 uL). The supernatant was obtained by centrifugation at 10,000 rpm for 20 min and then injected into HPLC for analysis. HPLC analysis was performed under the following conditions - mobile phase A: water with 0.1% TFA; B: acetonitrile with 0.1% TFA; 0-10 min: gradient elution, 2-95% B; 10-20min: isocratic elution, 95% B. The reaction was monitored using UV-vis absorbance at 600 nm. The peak was collected and was subjected to mass spectrometry analysis, respectively. The results further confirmed the observed peaks in HPLC trace are the enzymatic hydrolysis products.

**Synthesis of fluorescence probe**

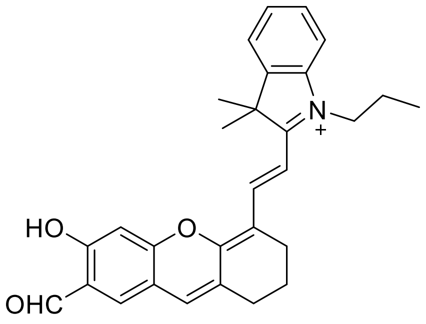


**2b**

[1]H NMR (500 MHz, $CDCl_3$) δ 10.04 (s, 1H), 8.66 (d, *J* = 15.1 Hz, 1H), 7.65 (s, 1H), 7.54 – 7.50 (m, 2H), 7.48 – 7.45 (m, 1H), 7.38 (d, *J* = 7.8 Hz, 1H), 7.08 (s, 1H), 6.98 (s, 1H), 6.59 (d, *J* = 15.1 Hz, 1H), 4.36 (t, *J* = 7.3 Hz, 2H), 2.72 (t, *J* = 6.5 Hz, 2H), 2.68 (t, *J* = 6.5 Hz, 2H), 1.99 – 1.92 (m, 4H), 1.81 (s, 6H), 1.07 (t, *J* = 7.5 Hz, 3H). [13]C NMR (126 MHz, $CDCl_3$) δ 193.55, 179.00, 165.14, 159.63, 157.99, 146.53, 142.37, 141.35, 131.81, 131.35, 129.49, 128.29, 128.10, 122.81, 119.81, 115.86, 115.35, 113.20, 106.01, 104.16, 51.34, 47.43, 29.24, 28.14, 23.97, 21.62, 20.30, 11.49. HRMS (ESI) Calcd. For $C_{29}H_{30}NO_3^+$ $[M]^+$: 440.2220; found: 440.2215.

Synthesis of **3a**

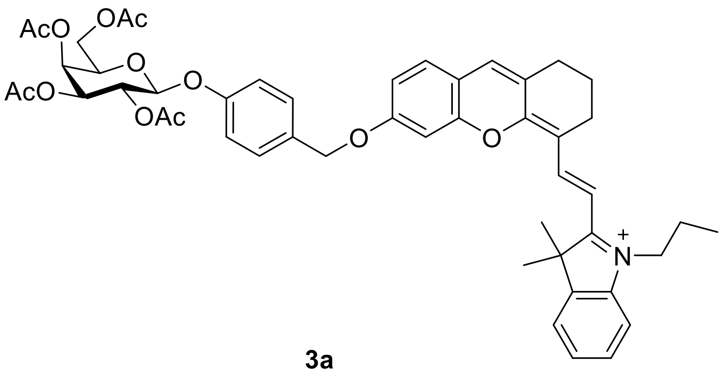

**3a**

To a solution of compound **2a** (27.0 mg, 0.05 mmol), $K_2CO_3$ (8 mg, 0.06 mmol) and NaI (14.9 mg, 0.1 mmol) in 2 mL of DMF, **1** (31 mg, 0.06 mmol) was added. Then the reaction mixture was stirred at room temperature in the dark for 3 days. HPLC was used to monitor reaction process. Once the reaction was completed. The reaction mixture was purified by HPLC to afford **3a** (18.0 mg, 37%).

$^1$H NMR (500 MHz, $CDCl_3$) δ 8.66 (d, *J* = 14.8 Hz, 1H), 7.51-7.47 (m, 2H), 7.43–7.37 (m, 4H), 7.30 (d, *J* = 7.9 Hz, 1H), 7.24 (s, 1H), 7.06 (d, *J* = 8.6 Hz, 2H), 6.97-7.96 (m, 2H), 6.35 (d, *J* = 14.8 Hz, 1H), 5.51 (dd, *J* = 10.4, 8.0 Hz, 1H), 5.47 (d, *J* = 3.2 Hz, 1H), 5.14- 5.12 (m, 3H), 5.09 (d, *J* = 7.9 Hz, 1H), 4.26 – 4.20 (m, 3H), 4.16 (dd, *J* = 11.2, 6.4 Hz, 1H), 4.09 (dd, *J* = *J* = 6.6 Hz, 1H), 2.75 (t, *J* = 5.5 Hz, 2H), 2.65 (t, *J* = 5.9 Hz, 2H), 2.19 (s, 3H), 2.08 (s, 3H), 2.06 (s, 3H) 2.02 (s, 3H), 1.96 – 1.93 (m, 4H), 1.79 (s, 6H), 1.07 (t, *J* = 7.4 Hz, 3H). $^{13}$C NMR (126 MHz, $CDCl_3$) δ 177.42, 170.62, 170.39, 170.31, 169.64, 162.34, 162.19, 157.19, 154.59, 145.96, 141.76, 141.55, 134.30, 130.60, 129.63, 129.33, 128.97, 127.54, 127.48, 122.72, 117.28, 116.13, 114.92, 113.83, 112.44, 103.36, 102.07, 99.65, 71.17, 70.92, 70.53, 68.72, 66.97, 61.42, 50.73, 46.84, 29.85, 29.24, 28.30, 24.10, 21.29, 20.87, 20.80, 20.73, 20.32, 11.53. HRMS (ESI) Calcd. For $C_{49}H_{54}NO_{12}^+$ $[M]^+$: 848.3641; found: 848.3608.

Synthesis of **3b**

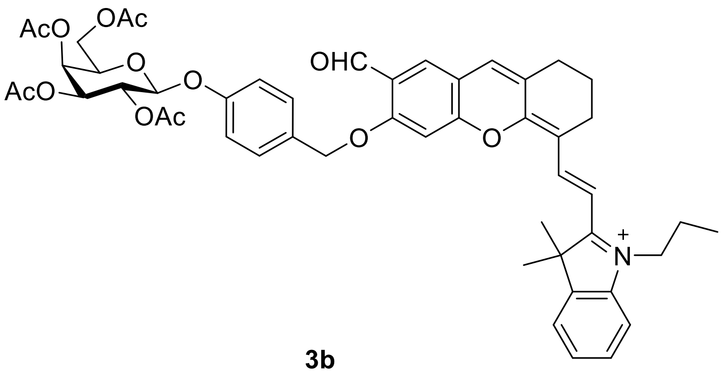


**3b**

This compound was synthesized in 32% yield with a similar procedure for **3a.**

$^1$H NMR (500 MHz, $CDCl_3$) δ 10.41 (s, 1H), 8.70 (d, *J* = 15.0 Hz, 1H), 7.84 (s, 1H), 7.58 (d, *J* = 7.2 Hz, 1H), 7.52 – 7.47 (m, 2H), 7.45 (d, *J* = 8.3 Hz, 2H), 7.36 (d, *J* = 7.7 Hz, 1H), 7.15 (s, 1H), 7.07 (s, 1H), 7.06 (d, *J* = 8.6 Hz, 2H), 6.49 (d, *J* = 15.0 Hz, 1H), 5.50 (dd, *J* = 10.3, 8.1 Hz, 1H), 5.47 (d, *J* = 3.2 Hz, 1H), 5.29 (s, 2H), 5.13 (dd, *J* = 10.5, 3.4 Hz, 1H), 5.10 (d, *J* = 7.9 Hz, 1H), 4.31 (t, *J* = 7.3 Hz, 2H), 4.23 (dd, *J* = 11.2, 6.9 Hz, 1H), 4.16 (dd, *J* = 11.2, 6.3 Hz, 1H), 4.10 (t, *J* = 6.6 Hz, 1H), 2.73 (t, *J* = 5.8 Hz, 2H), 2.66 (t, *J* = 5.8 Hz, 2H), 2.18 (s, 3H), 2.08 (s, 3H), 2.06 (s, 3H), 2.02 (s, 3H), 1.99 – 1.93 (m, 4H), 1.82 (s, 6H), 1.08 (t, *J* = 7.4 Hz, 3H). $^{13}$C NMR (126 MHz, $CDCl_3$) δ 187.90, 179.08, 170.58, 170.38, 170.26, 169.63, 163.73, 160.06, 158.09, 157.22, 146.62, 142.45, 141.23, 131.75, 130.16, 129.52, 129.39, 128.62, 128.40, 127.87, 123.06, 117.30, 115.69, 115.43, 112.96, 105.48, 101.00, 99.64, 71.30, 71.23, 70.96, 68.74, 67.01, 61.46, 51.40, 47.32, 29.39, 27.94, 24.19, 21.54, 20.88, 20.79, 20.78, 20.73, 20.24, 11.55. HRMS (ESI) Calcd. For $C_{50}H_{54}NO_{13}^+$ $[M]^+$: 876.3590; found: 876.3686.

Synthesis of **NIR-BG1**

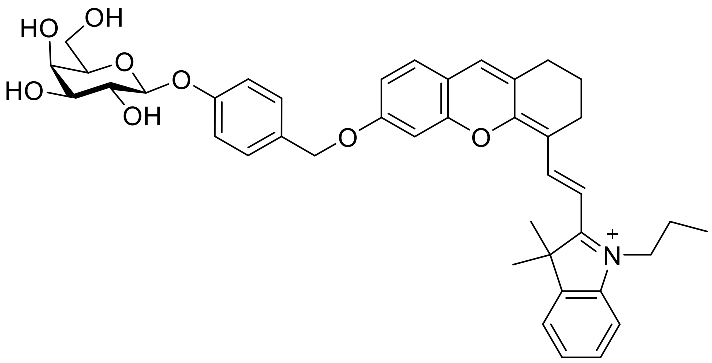

**NIR-BG1**

To a solution of **3a** (18 mg, 0.0187 mmol) in methanol (1 mL), NaOMe (0.43 µL) was added. The reaction was monitored with HPLC. After starting material was consumed, the crude product was purified with HPLC to afford NIR-BG1 in 68% yield.

$^{1}$H NMR (500 MHz, $CD_3OD$) δ 8.77 (d, *J* = 15.0 Hz, 1H), 7.68 (d, *J* = 7.5 Hz, 1H), 7.55-7.52 (m, 2H), 7.49-7.42 (m, 4H), 7.40 (s, 1H), 7.18-7.15 (m, 2H), 7.09 (d, *J* = 2.2 Hz, 1H), 7.04 (dd, *J* = 8.6, 2.4 Hz, 1H), 6.52 (d, *J* = 14.9 Hz, 1H), 5.21 (s, 2H), 4.88 (d, *J* = 8.0 Hz, 1H), 4.33 (t, *J* = 7.5 Hz, 2H), 3.90 (d, *J* = 3.0 Hz, 1H), 3.82-3.73 (m, 3H), 3.69 (dd, *J* = 6.8, 5.7 Hz, 1H), 3.58 (dd, *J* = 9.7, 3.4 Hz, 1H), 2.79 (t, *J* = 6.0 Hz, 2H), 2.72 (t, *J* = 6.0 Hz, 2H), 1.99-1.92 (m, 4H), 1.83 (s, 6H), 1.08 (t, *J* = 7.5 Hz, 3H). $^{13}$C NMR (126 MHz, $CD_3OD$) δ 179.29, 163.79, 163.13, 159.30, 155.82, 147.08, 143.50, 143.06, 135.17, 131.44, 130.29, 130.21, 130.06, 128.61, 128.40, 123.84, 117.94, 117.29, 115.64, 115.40, 113.92, 104.75, 102.91, 102.68, 77.06, 74.85, 72.24, 71.55, 70.20, 62.44, 52.02, 47.49, 30.05, 28.38, 25.03, 22.25, 21.65, 11.58. HRMS (ESI) Calcd. For $C_{41}H_{46}NO_8^+$ $[M]^+$: 680.3218; found: 680.3215.

Synthesis of **NIR-BG2**

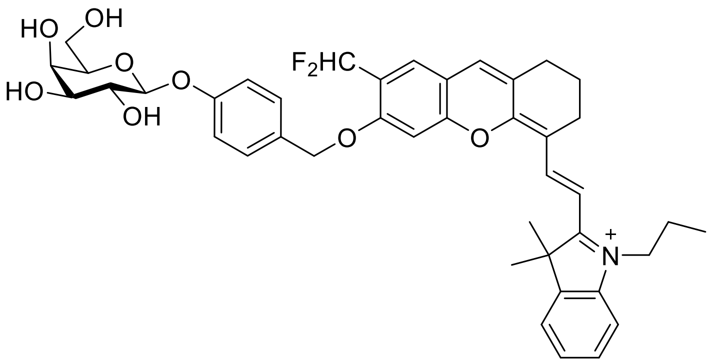

**NIR-BG2**

To a solution of **3b** (5.4 mg, 5.3 umol) in dichloromethane (1 mL), DAST (23 uL, 0.16 mmol) was added. The reaction was stirred at room temperature and monitored with HPLC. Once the starting material was consumed, the reaction was quenched with methanol for 1 h to consume remaining DAST. Next, the solvents were removed under reduced pressure, and the residue was dissolved in methanol. NaOMe was added to this mixture until the reaction was completed. Then the mixture was purified with HPLC to afford **NIR-BG2** in 70% yield over 2 steps.

$^1$H NMR (500 MHz, $CD_3OD$) δ 8.77 (d, $J$ = 15.0 Hz, 1H), 7.73–7.69 (m, 2H), 7.59 (d, $J$ = 15.0 Hz, 1H), 7.56 (t, $J$ = 7.6 Hz, 1H), 7.51 (t, $J$ = 7.4 Hz, 1H), 7.46 (d, $J$ = 8.5 Hz, 2H), 7.34 (s, 1H), 7.21– 7.18 (m, 3H), 7.00 (t, $J_{H\text{-}F}$ = 55.3 Hz, 1H), 6.60 (d, $J$ = 15.0 Hz, 1H), 5.33 (s, 2H), 4.88 (d, $J$ = 8.0 Hz, 1H), 4.38 (t, $J$ = 7.4 Hz, 2H), 3.89 (d, $J$ = 3.5 Hz, 1H), 3.82–3.72 (m, 3H), 3.69–3.67 (m, 1H), 3.56 (dd, $J$ = 9.7, 3.3 Hz, 1H), 2.78 (t, $J$ = 5.5 Hz, 2H), 2.72 (t, $J$ = 6.0 Hz, 2H), 1.99-1.93 (m, 4H), 1.86 (s, 6H), 1.09 (t, $J$ = 7.5 Hz, 3H). $^{13}$C NMR (126 MHz, $CD_3OD$) δ 180.14, 161.97, 160.69, 159.43, 156.82, 147.40, 143.79, 142.92, 133.44, 130.98, 130.31, 130.18, 129.66, 128.92, 126.71, 123.94, 118.11, 116.72, 115.88, 114.35, 112.55 ($J_{C\text{-}F}$ = 236.4 Hz), 106.06, 102.95, 101.84, 101.40, 77.10, 74.86, 72.25, 72.14, 70.22, 62.47, 52.37, 47.79, 30.11, 28.26, 28.24, 25.01, 22.40, 21.54, 11.56. HRMS (ESI) Calcd. For $C_{42}H_{46}F_2NO_8^+$ $[M]^+$: 730.3186; found: 730.3162.

[1] O. Redy-Keisar, E. Kisin-Finfer, S. Ferber, R. Satchi-Fainaro, D. Shabat, Synthesis and use of QCy7-derived modular probes for the detection and imaging of biologically relevant analytes, Nat Protoc, 9 (2014) 27-36.
[2] Z. Li, X.Y. He, Z. Wang, R.H. Yang, W. Shi, H.M. Ma, in vivo imaging and detection of nitroreductase in zebrafish by a new near-infrared fluorescence off-on probe, Biosens Bioelectron, 63 (2015) 112-116.

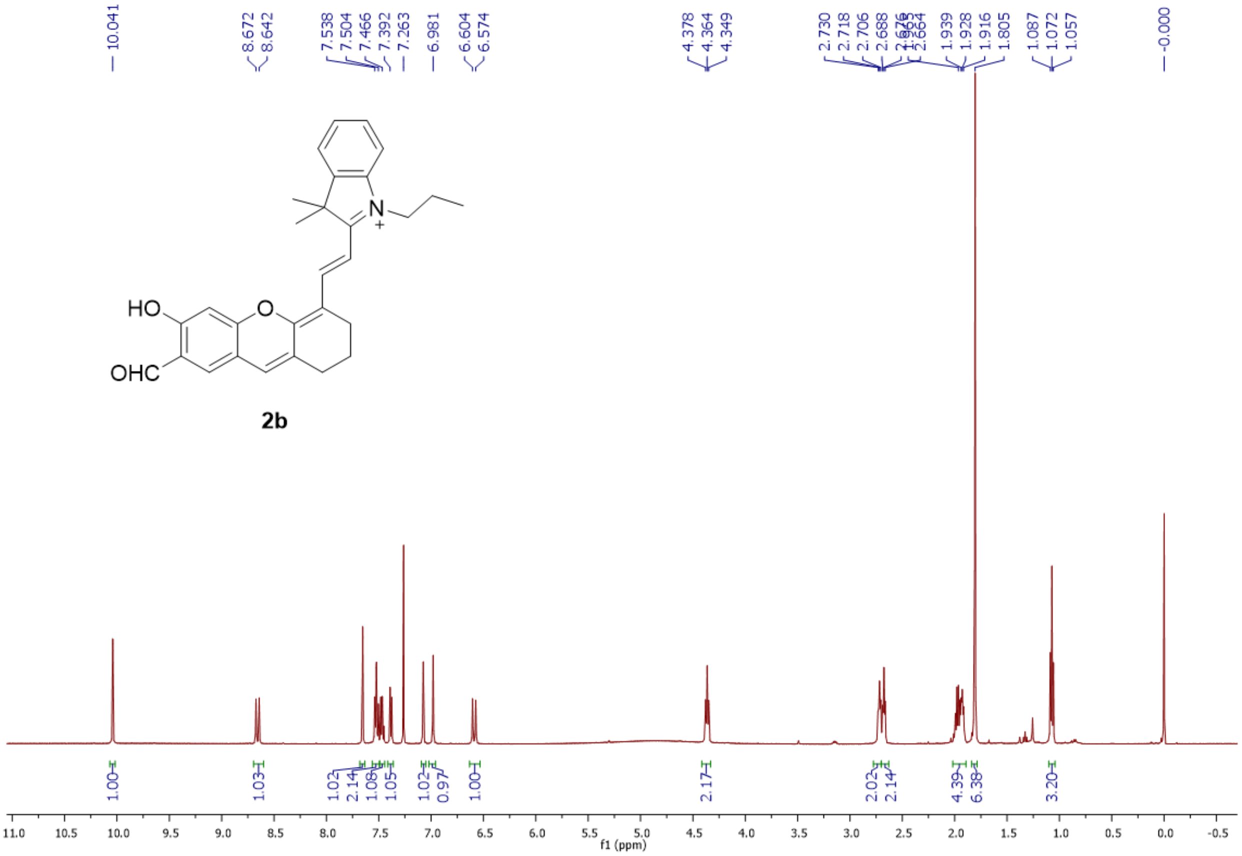


Fig. S1 $^{1}$H NMR spectrum of compound **2b** in $CDCl_3$.

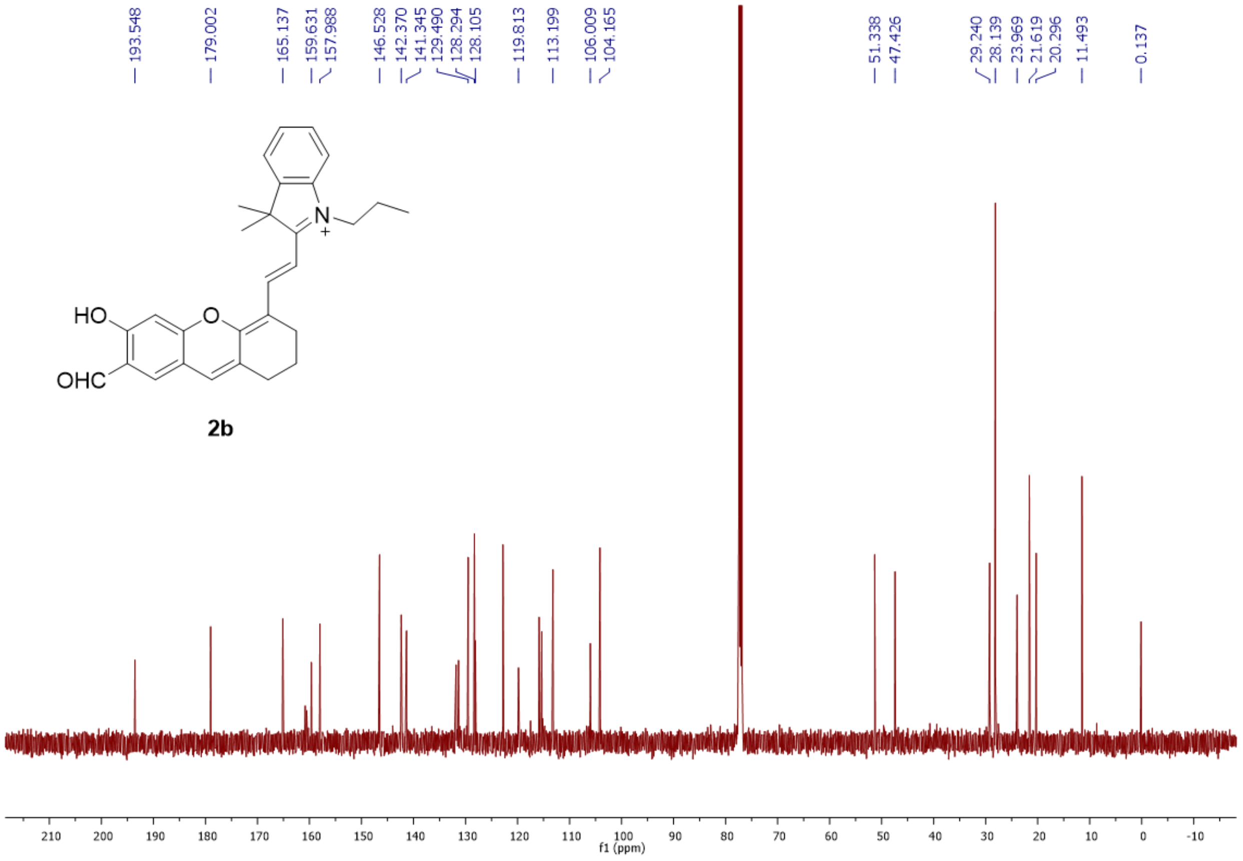


Fig. S2 $^{13}C$ NMR spectrum of compound **2b** in $CDCl_3$.

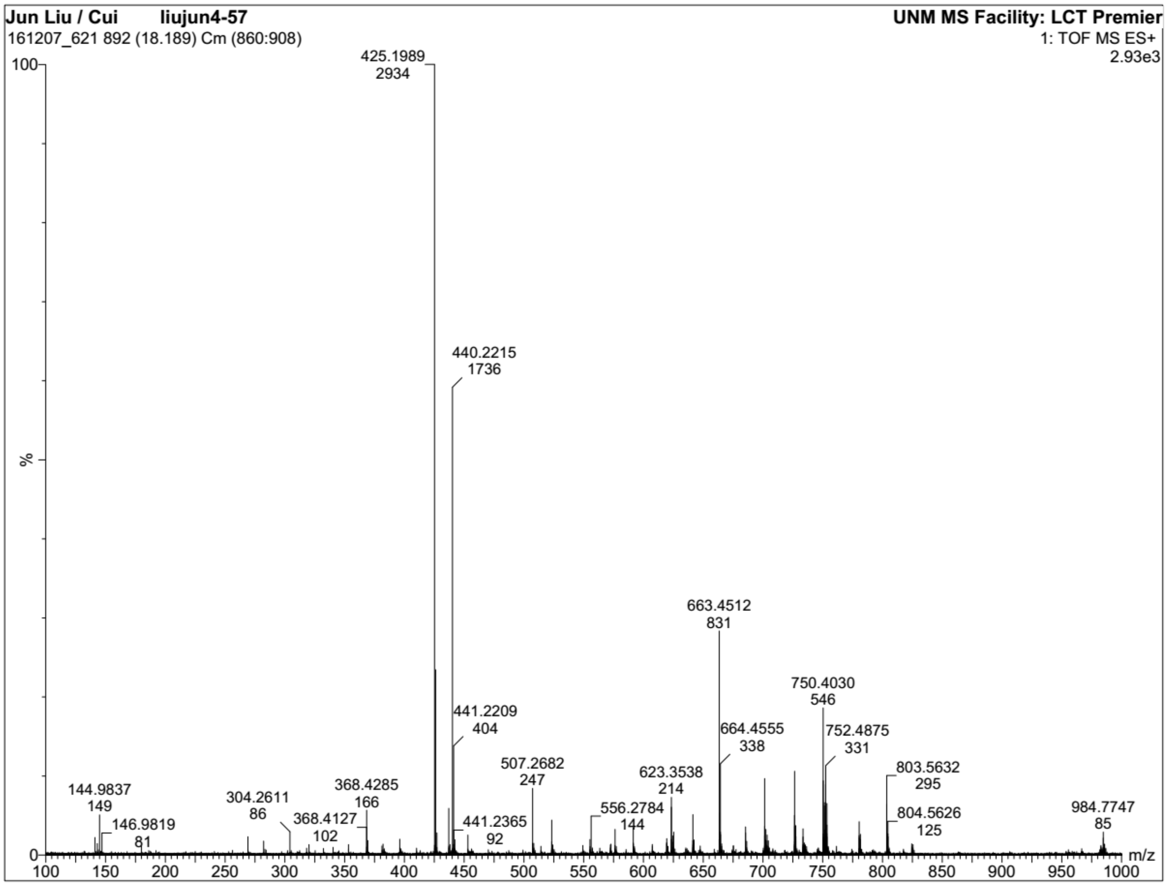


Fig. S3 ESI-MS spectra of **2b**

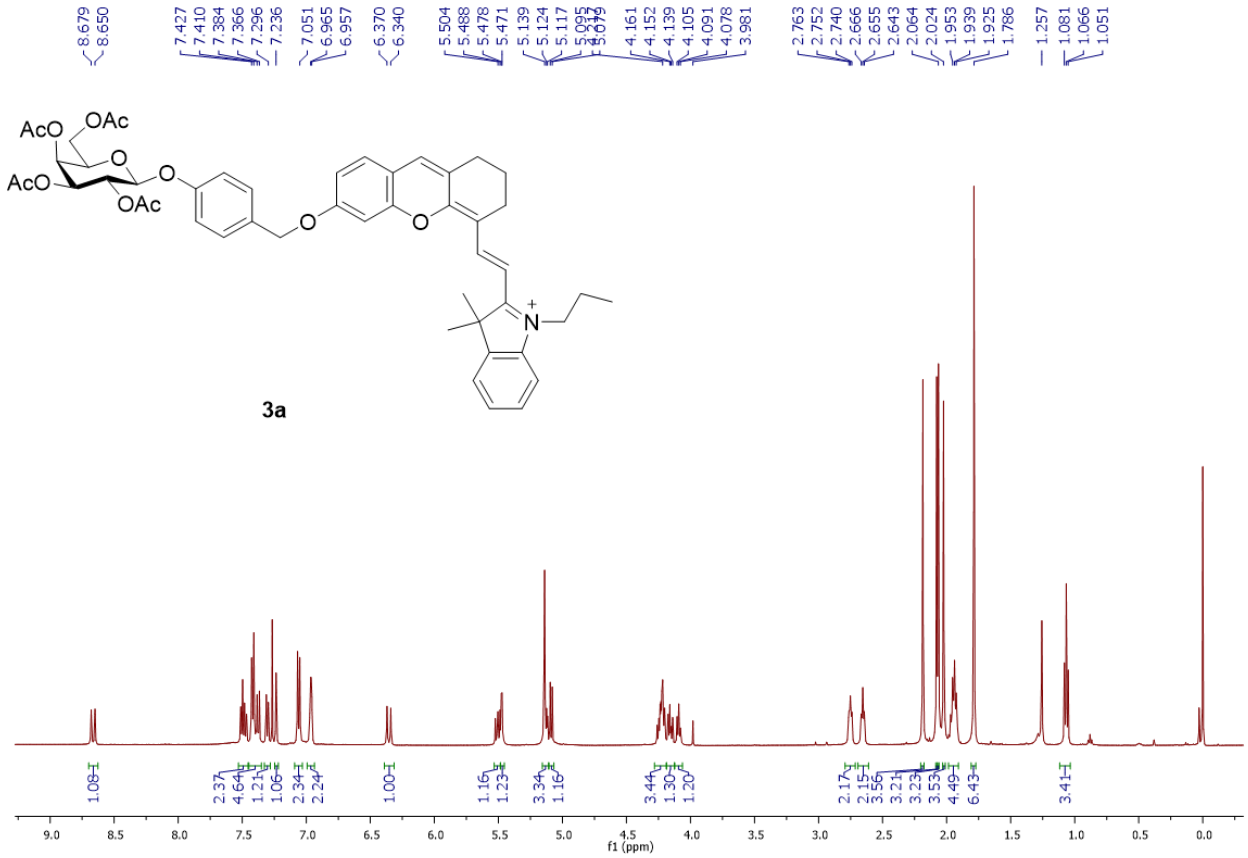

Fig. S4 $^{1}$H NMR spectrum of compound **3a** in $CDCl_3$.

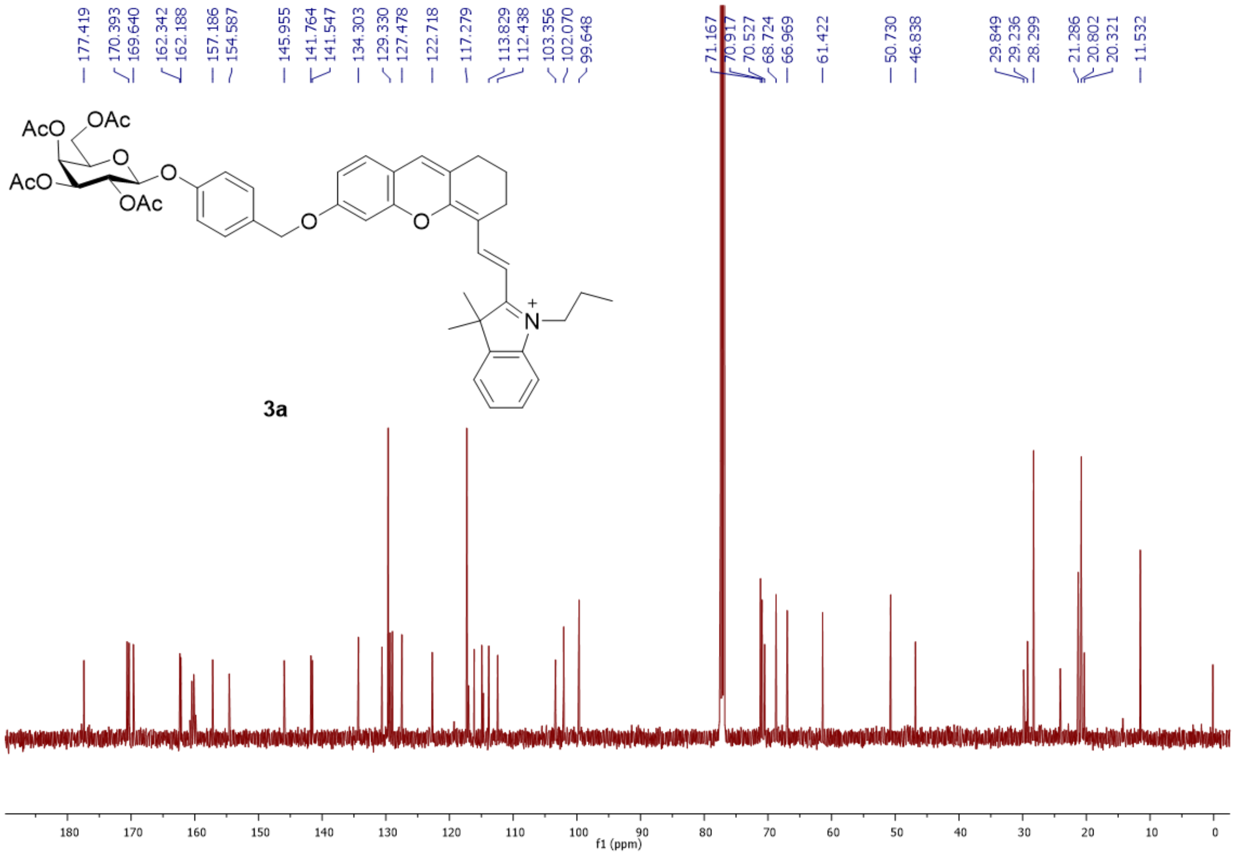


Fig. S5 $^{13}$C NMR spectrum of compound **3a** in $CDCl_3$.

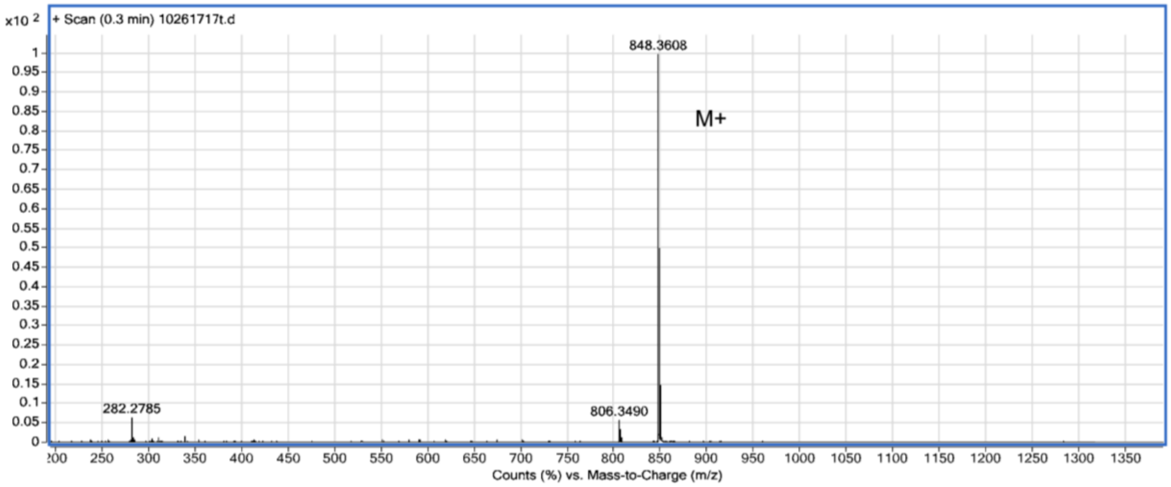


Fig. S6 ESI-MS spectra of **3a**

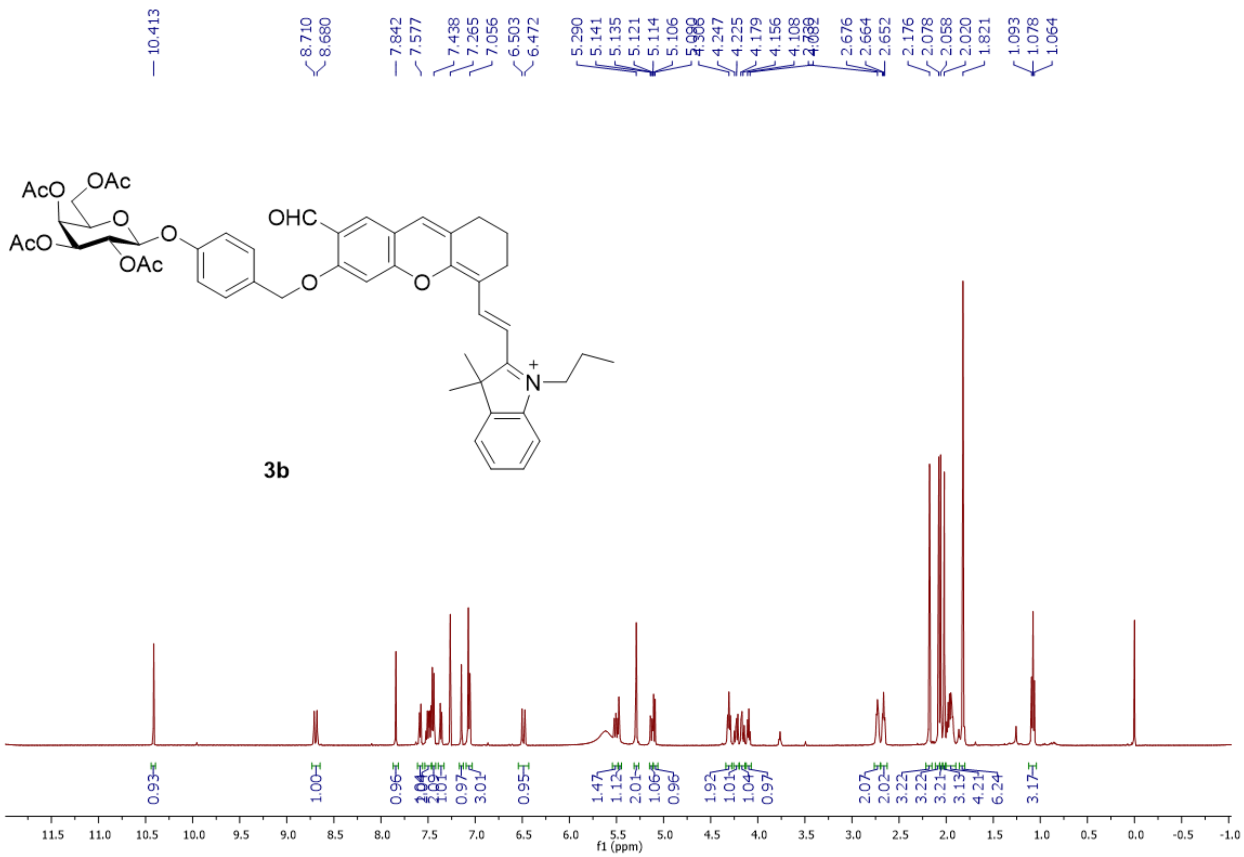

Fig. S7 $^{1}$H NMR spectrum of compound **3b** in $CDCl_3$.

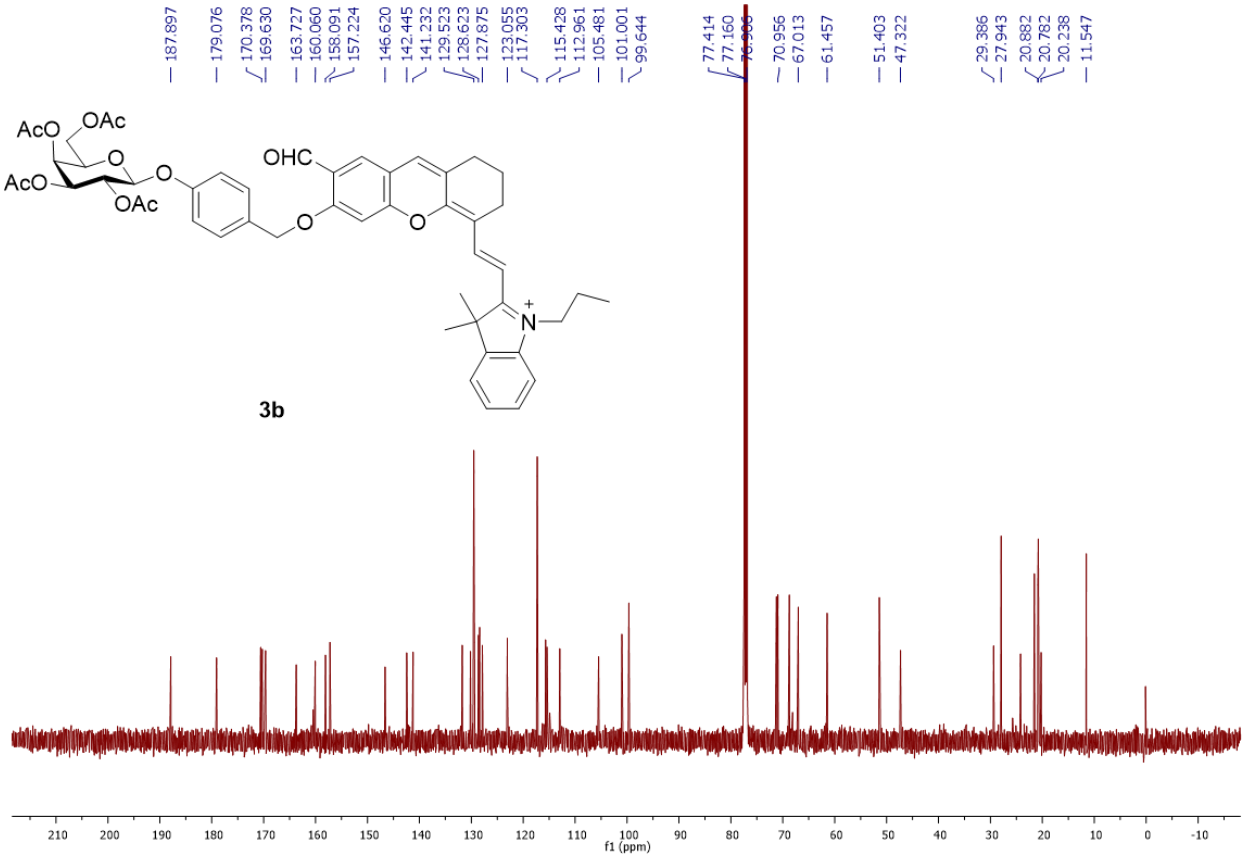


Fig. S8 $^{13}$C NMR spectrum of compound **3b** in $CDCl_3$.

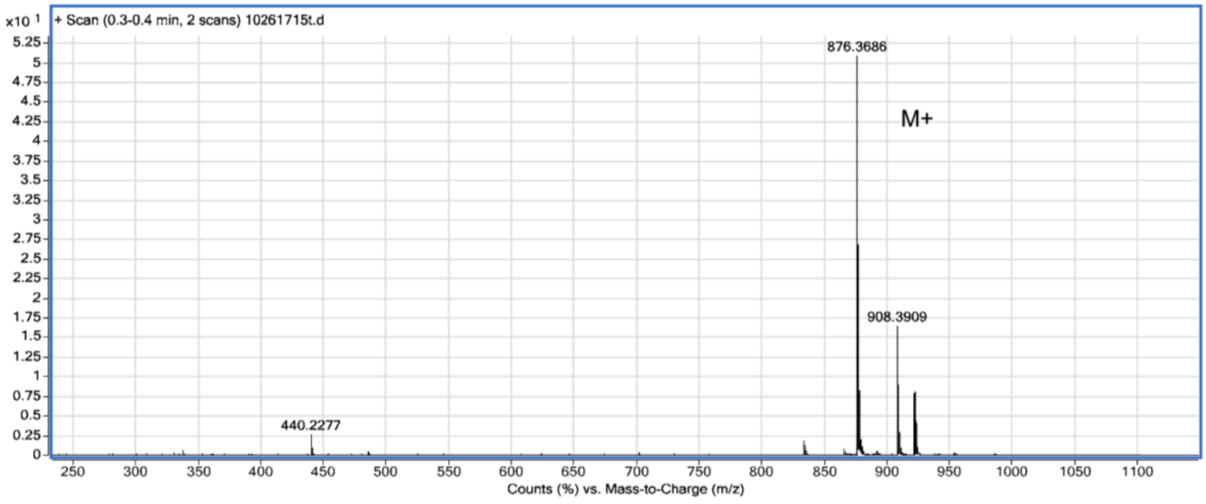


Fig. S9 ESI-MS spectra of **3b**

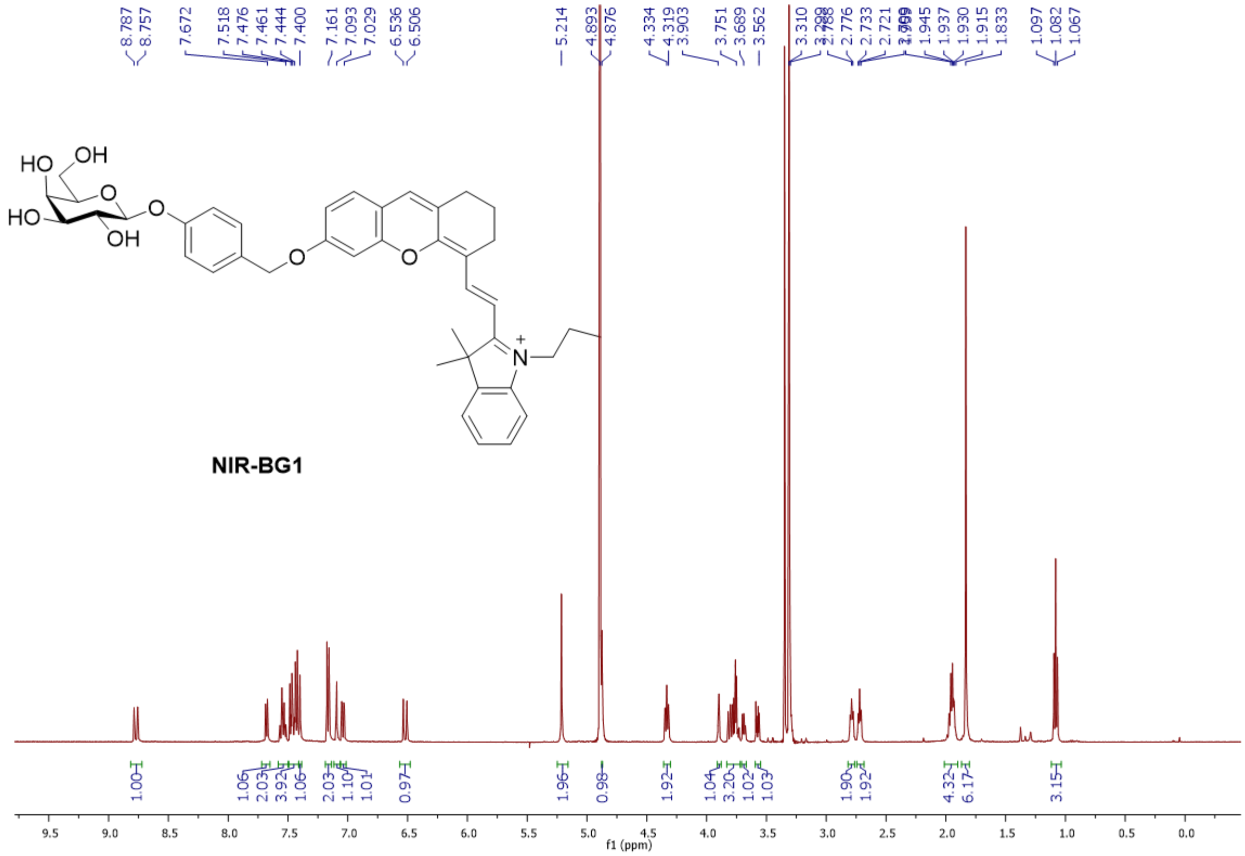

Fig. S10 $^{1}H$ NMR spectrum of compound **NIR-BG1** in $CD_3OD$.

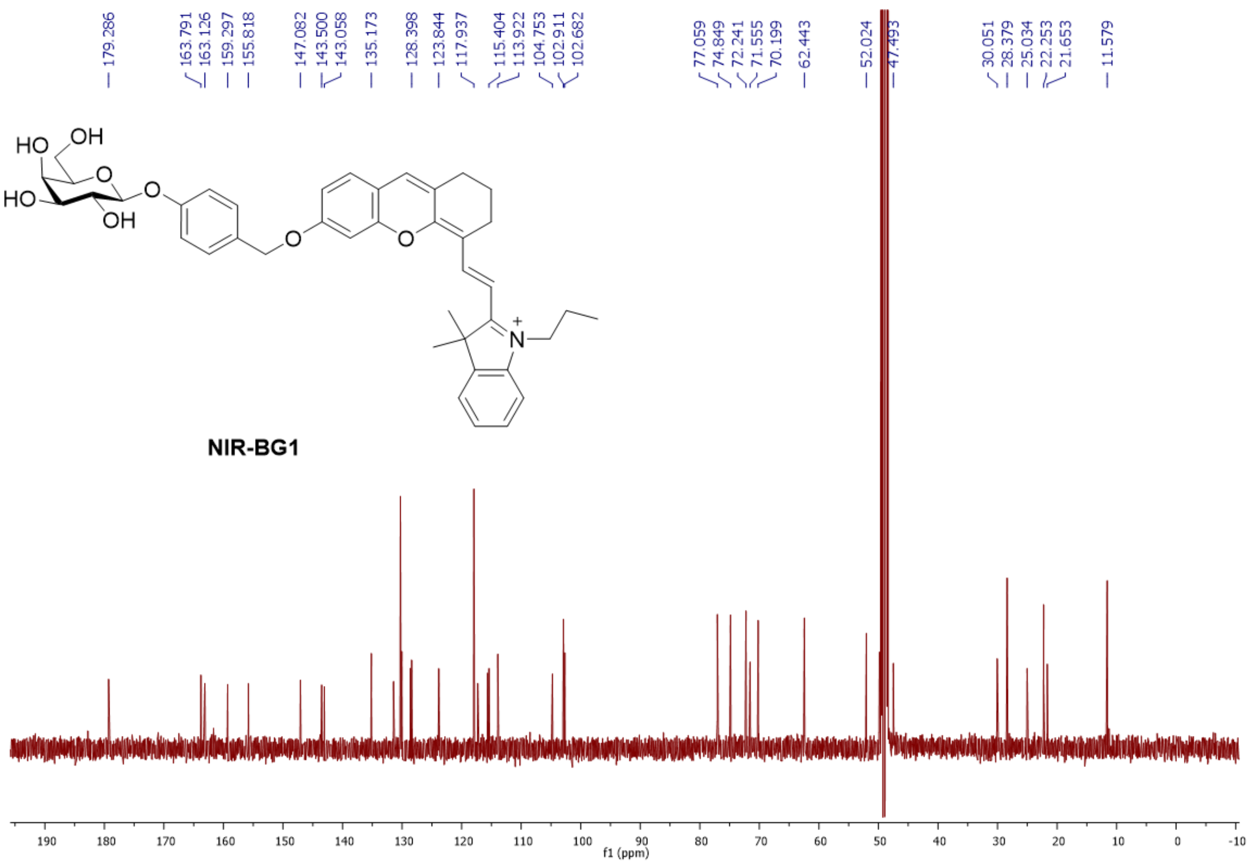


Fig. S11 $^{13}$C NMR spectrum of compound **NIR-BG1** in $CD_3OD$.

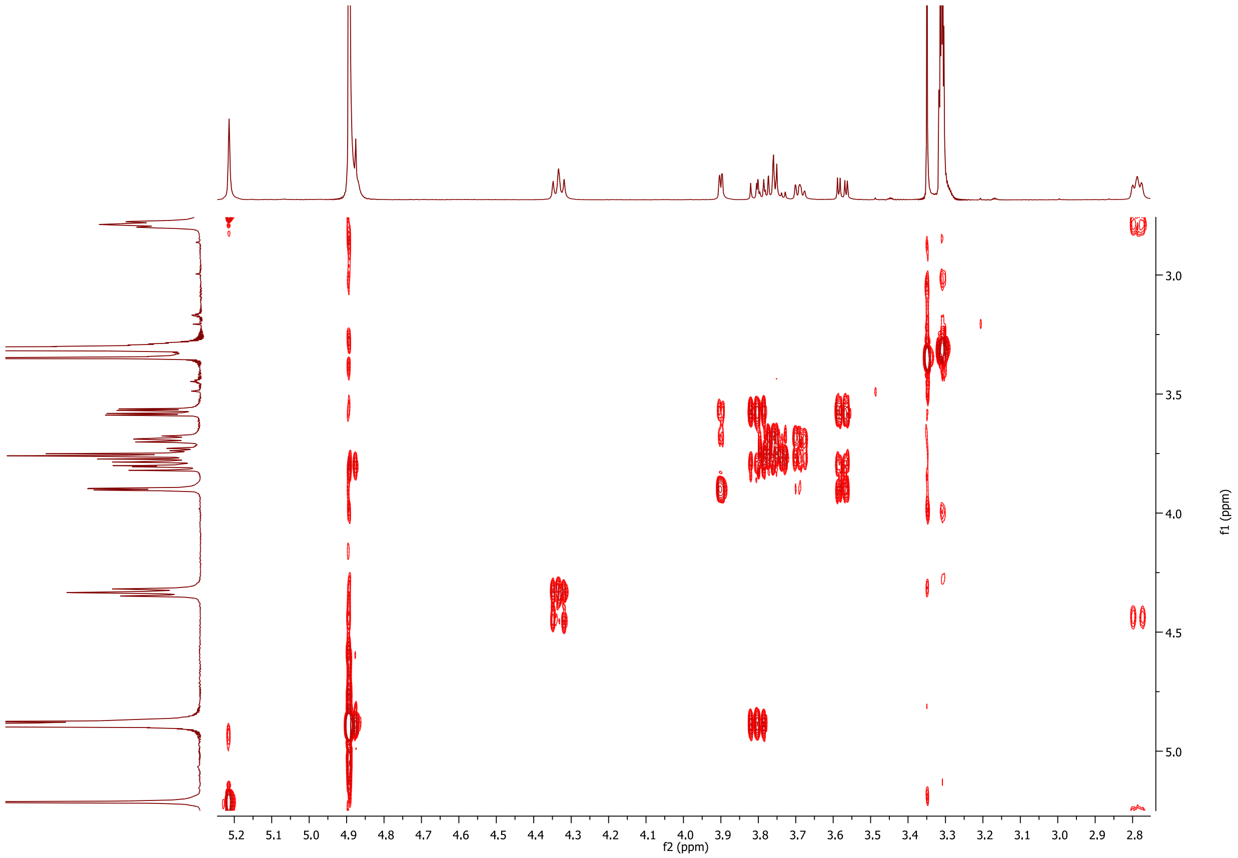


Fig. S12 [1]H-[1]H COSY spectrum of compound **NIR-BG1** in $CD_3OD$.

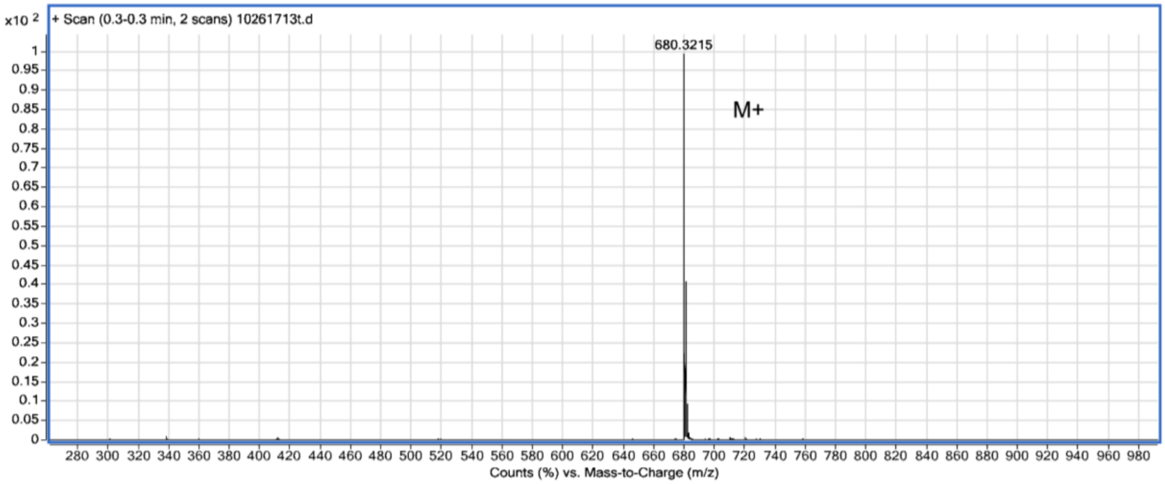

Fig. S13 ESI-MS spectra of **NIR-BG1**

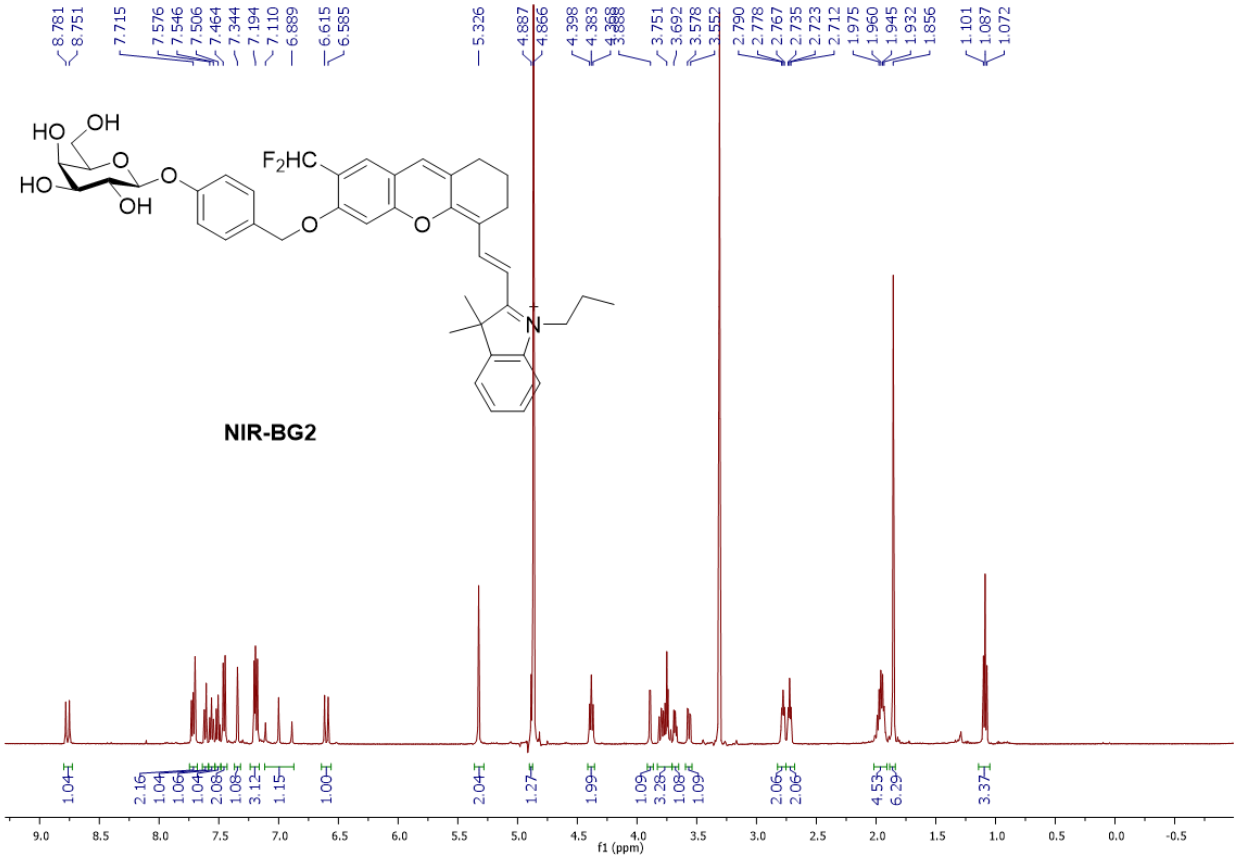


Fig. S14 $^{1}H$ NMR spectrum of compound **NIR-BG2** in $CD_3OD$.

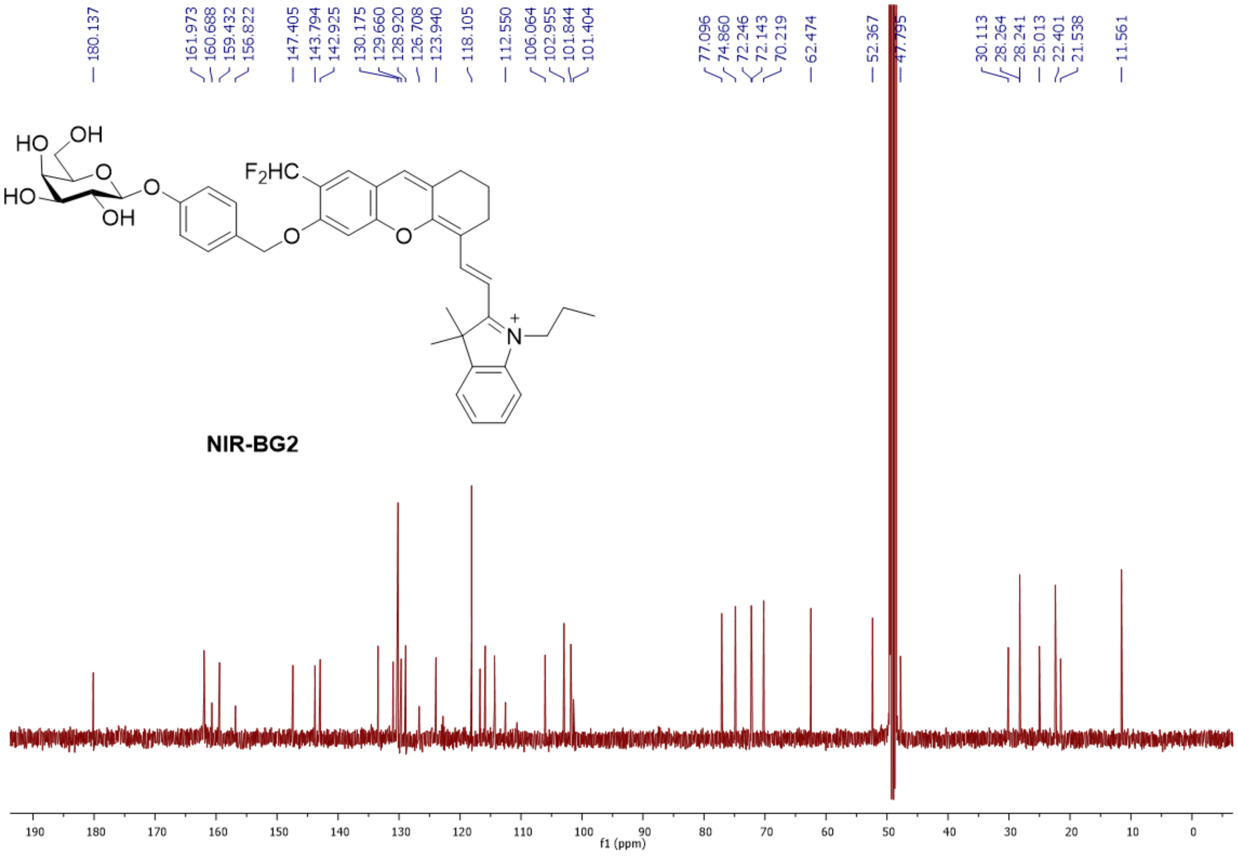


Fig. S15 $^{13}C$ NMR spectrum of compound **NIR-BG2** in $CD_3OD$.

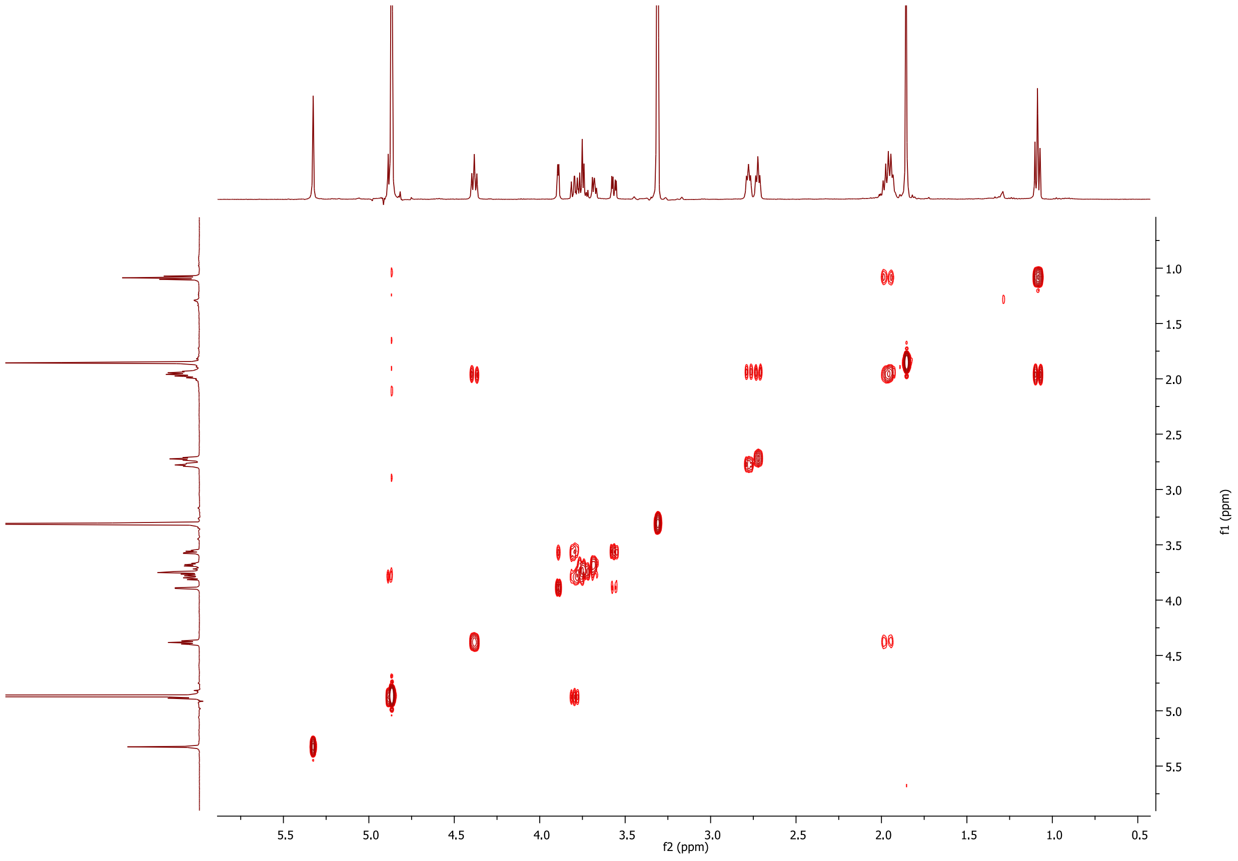


Fig. S16 $^1H$-$^1H$ COSY spectrum of compound **NIR-BG2** in $CD_3OD$.

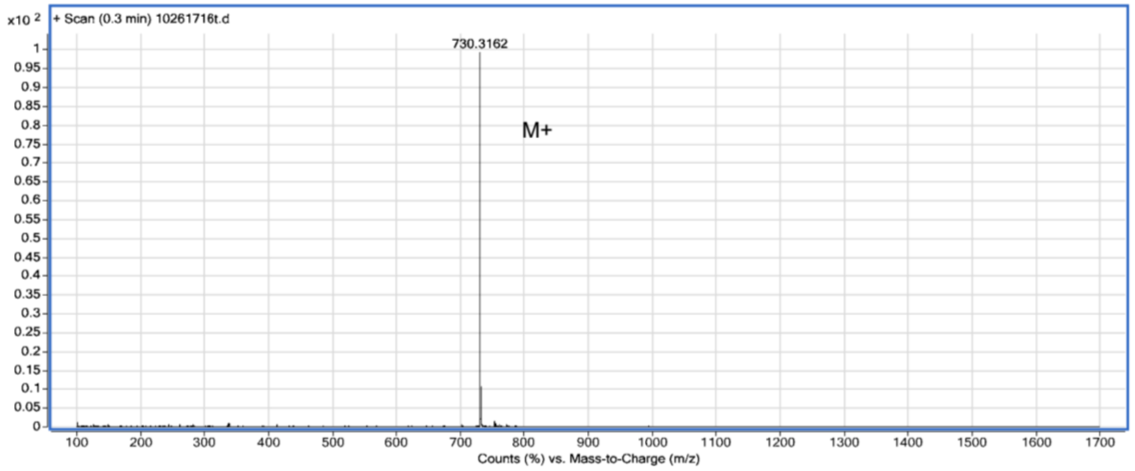


Fig. S17 ESI-MS spectra of **NIR-BG2**
